# A regulatory network of Sox and Six transcription factors initiate a cell fate transformation during hearing regeneration in adult zebrafish

**DOI:** 10.1101/2022.02.09.479753

**Authors:** Erin Jimenez, Claire C. Slevin, Wei Song, Zelin Chen, Stephen C. Frederickson, Derek Gildea, Weiwei Wu, Abdel G. Elkahloun, Ivan Ovcharenko, Shawn M. Burgess

## Abstract

Using adult zebrafish inner ears as a model for sensorineural regeneration, we performed a targeted ablation of the mechanosensory receptors in the utricle and saccule and characterized the single-cell epigenome and transcriptome at consecutive time-points following hair cell ablation. Using deep learning on the regeneration-induced open chromatin sequences, we were able to identify unique, cell-specific transcription factor (TF) motif patterns enriched in the raw data. We correlated enhancer activity with gene expression to identify gene regulatory networks. A clear pattern of overlapping Sox- and Six-family transcription factor gene expression and binding motifs was detected, suggesting a combinatorial program of TFs driving regeneration and cell identity. Pseudo-time analysis of single-cell transcriptomic data demonstrated that the support cells within the sensory epithelium changed cell identity to a more pluripotent “progenitor” cell population that could either differentiate into hair cells or return to a support cell identity. We showed that *sox2* becomes enriched in the progenitor cells and is reduced again when the cells differentiate in either direction. Analysis of the scATAC-seq data identified a 2.6 kb DNA sequence element upstream of the *sox2* promoter that dynamically changed in accessibility during hair cell regeneration. When deleted, the upstream regulator of *sox2* showed a dominant phenotype that resulted in a hair cell regeneration-specific deficit in both the lateral line and adult inner ear.

**HIGHLIGHTS:**

Integrated scRNA-seq and scATAC-seq of the adult zebrafish inner ear during hair cell regeneration.
Support cells transition to an intermediate cell type that can differentiate to hair cells. Coordinated expression of Sox and Six transcription factors are key to hearing regeneration.
Sox transcription factors trigger the regeneration response in the support cells while Sox and Six factors cooperate during hair cell differentiation.
Deletion of an upstream enhancer that controls the timing of *sox2* expression causes regeneration-specific defects in hearing regeneration.

## INTRODUCTION

The capacity to regenerate tissues after injury unevenly manifests across the vertebrate lineage (Sánchez Alvarado, 2003). Why some vertebrate animals are able to regenerate multiple tissues and others cannot continues to be a fundamental question in regenerative biology. In mammals, consistent cellular renewal is limited to certain cell types such as skin, gut, and blood, while major tissue regeneration is even further restricted to a small number of organs such as the liver. Many tissues in mammals fail to induce any regenerative response after injury including the inner ear sensory epithelium. Damage to the mammalian inner ear sensory epithelium is irreversible and results in permanent hearing loss or vestibular defects. Interestingly, this is a feature that sets mammals apart from most other vertebrates who can continually produce new hair cells throughout their lifetimes and/or can regenerate them in response to trauma. Investigations of the genes involved in hair cell regeneration have shown that the key gene players in development can also have important roles in regeneration (Groves et al., 2013). However the regeneration phenomenon is distinct from development (Pei et al., 2018) and recent genome-wide analyses suggest that while regeneration programs may target many of the same genes, they may do so through distinct regulatory sequences (Kang et al., 2016; Suzuki and Ochi, 2020); reviewed in (Rodriguez and Kang, 2020; Yang et al., 2019).

Enhancer regulatory elements have been shown to be critical in the control of development (Ryan and Farley, 2020), and now several groups have made connections between enhancer regulation and various tissue regeneration programs. Injury-responsive or regeneration-associated enhancers that direct gene expression in injured tissues have been identified in the regenerating heart and fin of zebrafish (Begeman et al., 2020; Goldman et al., 2017; Harris et al., 2016; Kang et al., 2016; Lee et al., 2020; Rodriguez and Kang, 2020; Soukup et al., 2019; Suzuki et al., 2019; Thompson et al., 2020; Vizcaya-Molina et al., 2018; Wang et al., 2006). Comparative epigenomic profiling and single-cell genomics experiments have revealed species specific and evolutionarily conserved genomic responses to regeneration in fish (Goldman et al., 2017; Kang et al., 2016; Wang et al., 2020). Multiple “tissue regeneration enhancer elements” (TREEs) that govern regeneration-dependent gene expression in adult zebrafish and killifish fins and hearts have been identified. However, ablation of enhancers in these studies showed that these injury responsive candidates are generally not essential for normal regeneration.

Here, we profiled genome-wide changes in chromatin accessibility (scATAC-seq) and gene expression (scRNA-seq) during regeneration of the zebrafish inner ear at single-cell resolution to identify distinct phases of inner ear hair cell regeneration and to correlate cell-state changes with key gene regulatory networks. We showed the support cells transitioned into a “progenitor-like” state that could differentiate into new hair cells or return to support cells. We also identified a key regulator of *sox2* expression that when deleted, the hair cells developed normally, but hair cell regeneration after injury was significantly disrupted.

## RESULTS

### Single-cell RNA sequencing identifies different cell populations in the inner ear

For the hair cell ablation studies, we used the Tg(*myo6b*:DTR) transgenic zebrafish which permits conditional and selective ablation of hair cells in the adult zebrafish (Jimenez et al., 2021). Treatment of adult Tg(*myo6b*:DTR) zebrafish with a single 0.05 ng/µL injection of diphtheria toxin (DT) leads to complete hair cell loss of auditory (saccule) and vestibular (utricle) hair cells 5 days-post injection leaving a population of differentiated *sox2* positive supporting cells undamaged (Jimenez et al., 2021) (Figure 1A). To elucidate the transcriptional regulatory network that controls tissue regeneration, we characterized the gene expression profiles (scRNA-seq) and the map of accessible regions (scATAC-seq) associated with the response to hair cell ablation in adult zebrafish sensory epithelia. We dissected out saccules and utricles at three time points: Days 4, 5 and 7 post-DT, and each were processed separately in both scRNA-seq and scATAC-seq experiments. The first time point (Day 4) corresponds to maximal hair cell clearance after the apoptotic program has been initiated. The second time point (Day 5) corresponds to when hair cells remain absent but regeneration has been clearly initiated. Finally, the third selected time point (Day 7) corresponds to when hair cells begin to repopulate sensory epithelia (Figure 1B, C). Untreated Tg(*myo6b*:DTR) transgenic zebrafish and wild-type DT injected zebrafish lacking the Tg(*myo6b*:DTR) transgene were used as non-regenerating controls. Subsequent single-cell analysis entailed pairwise comparisons between controls and regenerating sensory epithelia at each time point.

**Figure 1.**
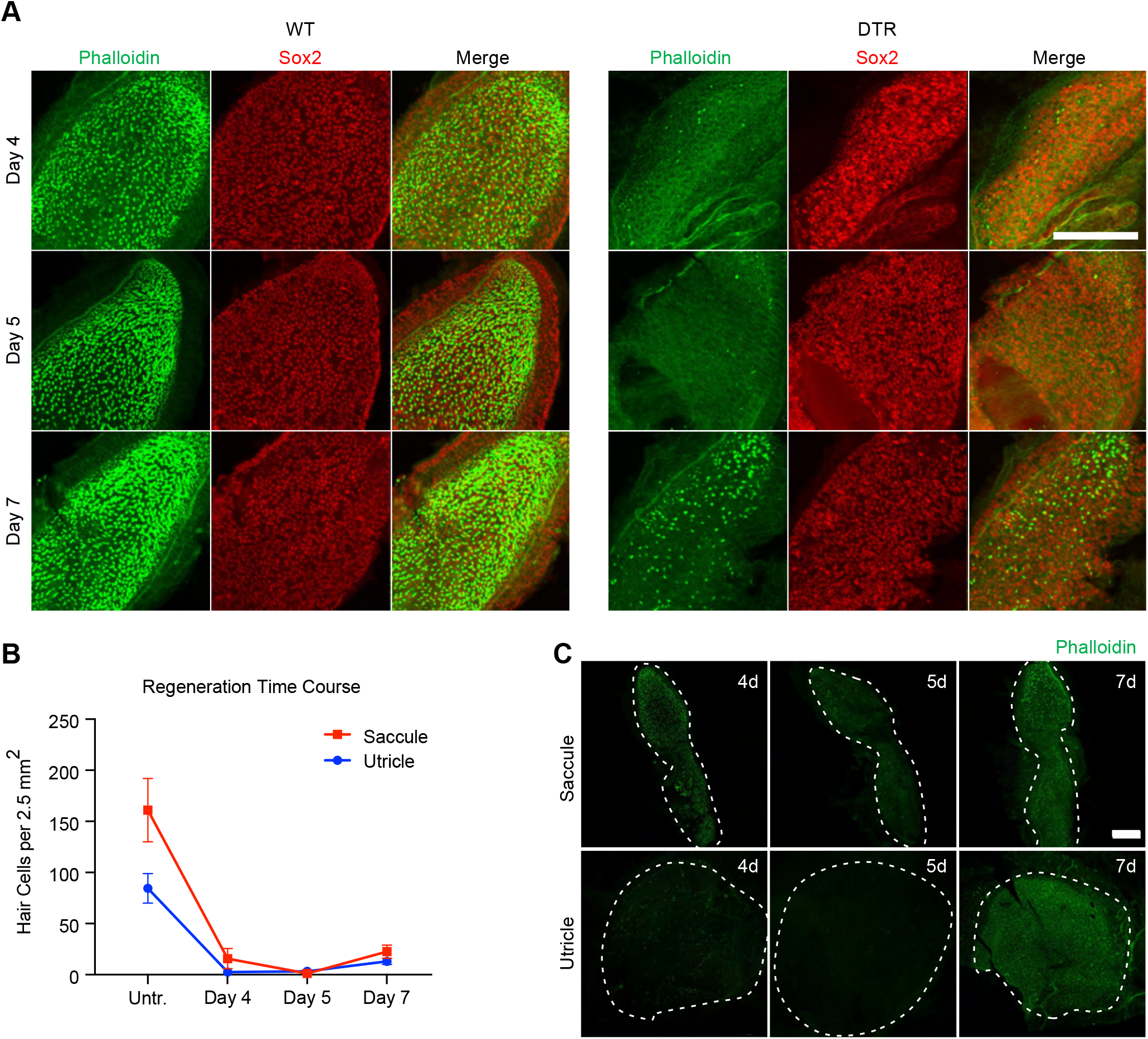
Vestibular and auditory hair cell regeneration following targeted ablation with diphtheria toxin in zebrafish. (A) Saccule isolated from wild-type (WT) and Tg(*myo6b*:hDTR) (DTR) zebrafish following DT injection on days 4, 5, and 7. Phalloidin staining (green) labels hair cell stereocilia and Sox2 immunostaining (red) labels supporting cells. Scale bar 100 µm. (B) Quantification of phalloidin positive hair cell number in saccule and utricle of untreated Tg(*myo6b*:hDTR) zebrafish and DT injected Tg(*myo6b*:hDTR) zebrafish on days 4, 5, and 7. Error bars demonstrate the mean ± SEM. (C) Saccule (top) and utricle (bottom) isolated from DT injected Tg(*myo6b*:hDTR) zebrafish on days 4, 5, and 7. Scale bar 100 µm.

We generated twelve transcriptomic profiles on inner ear tissue using the 10X Chromium system for droplet-based single-cell 3’ RNA sequencing (scRNA-seq) and quantified each data set using the Cell Ranger 6.0.0 pipeline (10x Genomics). We used the Seurat package on the scRNA-seq data to filter and process the resulting data. Cells that had unique feature counts over 1000 or less than 200 and over 5% mitochondrial counts were filtered. After filtering, we integrated datasets from all twelve samples for cell type clustering (Becht et al., 2018; Stuart et al., 2019). Samples were assembled into an aggregate or “unified transcriptomic atlas”. Unsupervised clustering partitioned 66,296 inner ear sensory epithelial cells (saccule and utricle combined) into discrete scRNA-seq cell populations (Figure 2A; Table S6). We assigned cell type identities to each of these clusters based on the top differentially expressed genes associated with each cluster and the known expression of these genes based on the literature using zebrafish transcriptome data from inner ear cells (Barta et al., 2018) and single-cell transcriptome data from the larval lateral line (Baek et al., 2021; Lush et al., 2019).

**Figure 2.**
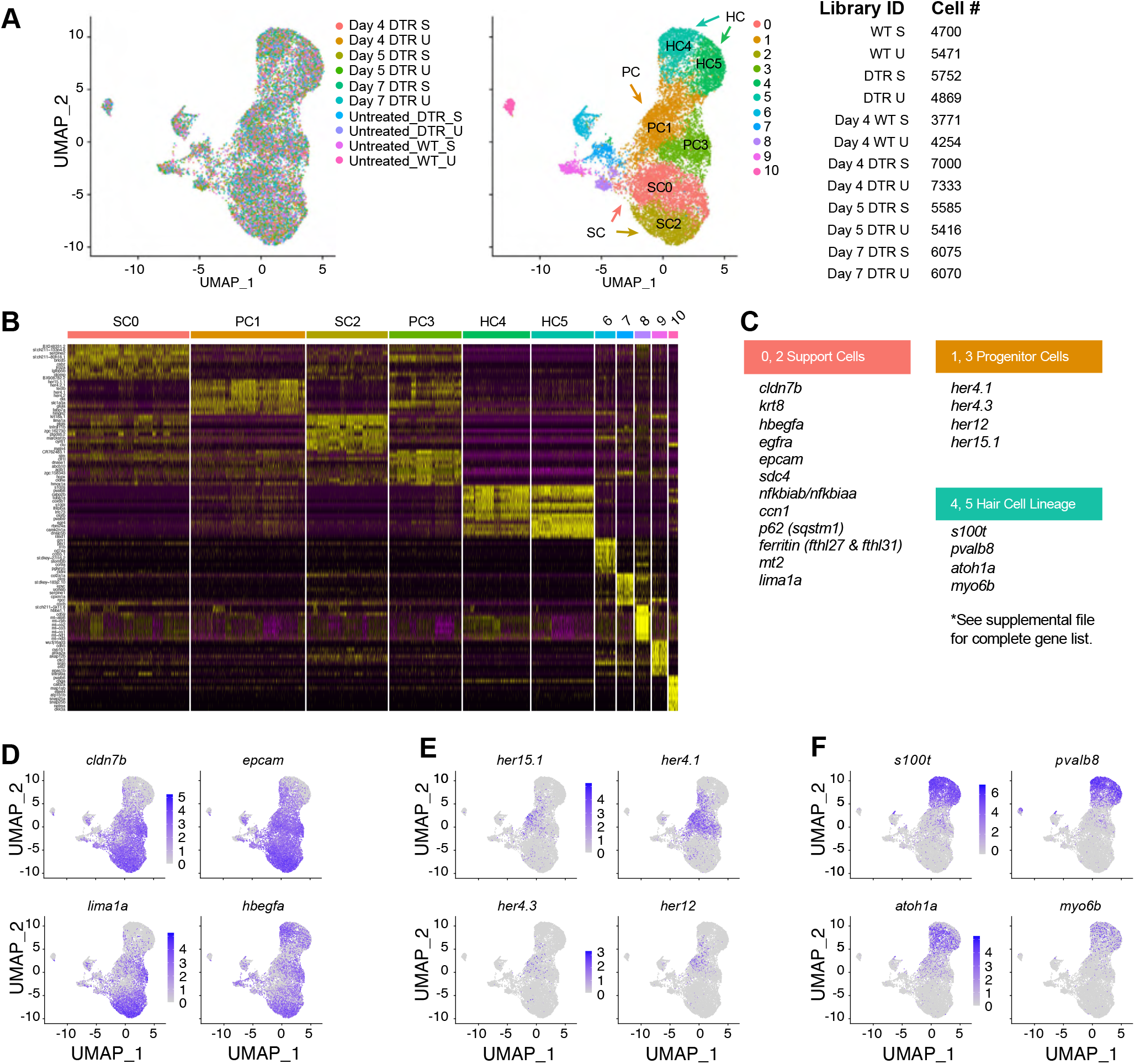
scRNA-seq of the auditory and vestibular epithelia identifies basal gene expression. (A) Cell type clustering assignments for inner ear sensory epithelia (saccule and utricle aggregate) collected from each timepoint including untreated controls. Uniform Manifold Approximation and Projection (UMAP) axes were calculated. Cluster 0 to 5 consist of cells that contribute to the sensory epithelium: Support Cells (SC), Progenitor Cells (PC), and Hair Cells (HC). Cell counts for each of the samples collected for scRNA-seq. (B) Heatmap showing the relative expression levels of the top 10 differentially expressed genes (y-axis) from each cluster (x-axis) of sensory epithelia sorted by highest fold change. The cell population identity assigned to each cluster is indicated above each column and colors correspond to clusters in B. (C) Table of top significant marker genes that distinguish the cell clusters. (D) UMAP plots of known marker genes overrepresented in support cells, (E) progenitor cells, (F) and hair cells.

For each cell population we identified genes specifically expressed or highly enriched (Figure 2B). We controlled for global gene expression responses caused by dissection, dissociation and other manipulations by selecting genes that were variable between clusters and/or between samples. We assigned cell types that directly contributed to new hair cells as: support cells (SC), progenitor cells (PC), young (yHC) and mature hair cells (mHC). Based on marker gene expression, we detected 2 populations of support cells (clusters SC0 and SC2) which showed significant enrichment for the markers *cldn7b*, *epcam*, *lima1a*, and *hbegfa* (Figure 2D). Support cell marker genes were enriched in clusters furthest from hair cell specific clusters, were reduced in intermediate cell clusters, and significantly more reduced in the hair cell clusters. We observed a distinct intermediate population of progenitor cells which expressed *her4.1*, *her15.1*, and *dla* (Figure 2E). The progenitor cells (PC) belong to clusters 1 and 3. This population of cells included both cycling (PC1) and non-cycling cells (PC3). Clusters 4 (yHC) and 5 (mHC) encompassed the hair cell (HC) lineage and expressed the *atoh1a*, *s100t*, and *pvalb8* genes (Figure 2F). No other cell cluster expressed hair cell lineage genes. Cells in all other clusters represented multiple non-sensory cell populations, such as immune cells, that do not directly contribute to the sensory epithelium and these were excluded from downstream analyses regardless if they responded to hair cell injury. Having established the different support, progenitor, and hair cell populations of the inner ear allowed us to interrogate the dynamic expression pattern of specific genes of interest in non-regenerating and regenerating inner ear tissues.

### Adult inner ear cells are transcriptionally distinct from larval lateral line cells

The zebrafish lateral line has frequently been used as a model for hair cell regeneration, so we examined how much overlap the two organs showed in terms of the regenerative program. Adult inner ear sensory epithelia demonstrated distinct differences from previously FACS-purified neuromasts (Figure S1; Table S1). There was strong separation of cell location and limited mixing of cells within clusters (Figure S1A). There is a clear geographic separation of cell types (Figure S1B). Figure S1C highlights the distinct gene expression profile of support cells, progenitor cells, and hair cells of adult inner ear. The differential distribution of the genes *s100t*, *atoh1a*, *cldn7b* were examined. The results are consistent with significant transcriptional differences in cells between neuromasts and adult inner ears (Figure S1D). The differences in transcriptome support the transcriptional distinctiveness of adult inner ear sensory epithelia from their larval lateral line counterparts suggesting a value to examining core conserved pathways as well as tissue specific ones between the two organs.

### Similarities between gene expression in the saccule and utricle of the zebrafish inner ear

The zebrafish auditory (saccule) and vestibular (utricle) organs are distinct sub-organs that share morphological and physiological aspects. The utricle in fish primarily functions as a gravitation sensor but has also shown auditory potential (Popper, 2011; Schulz-Mirbach and Ladich, 2016). We compared saccule and utricle scRNA-seq experiments to identify cell types that are unique or shared between these two otolith organs, to obtain conserved cell type markers and to also find cell-type specific responses in regenerating sensory epithelia (Figure S2; Table S2 and S3) (Butler et al., 2018). First, we examined the cellular diversity between untreated or homeostatic saccule and utricle (Figure S2A). Despite different morphological profiles (Bever and Fekete, 2002; Schulz-Mirbach and Ladich, 2016), saccule and utricle sensory and non-sensory cells are transcriptionally very similar and cluster together (Figure S2A). We tested if there were significant differences between regenerating saccule and utricle scRNA-seq experiments to determine if there was a divergence in regeneration of the auditory and vestibular sensory epithelia (Figure S2B). We first looked broadly at changes in gene expression between regenerating saccule and utricle cell types and found very few genes that differ (Figure S2C; Table S4). We then identified shared cell states between the regenerating saccule and utricle through integrated clustering analysis and found that gene expression changes are largely conserved between datasets for the three major cell populations that contribute to hair cell regeneration: SC, PC and HC (Figure S2C; Table S5). In agreement with Yao et al. (Yao et al., 2020), we observed elevated expression levels of *wnt11r*, *sema3e*, *otol1a*, and *nr2f1* in the saccule and identified that these genes are expressed in the support cell clusters specifically. During regeneration, these genes are upregulated only in the saccule. The gene *otofa* for example has higher expression in the utricle compared to the saccule, yet both tissues express this gene during hair cell differentiation (Figure S2). Since the regeneration programs between organs were so similar, we decided to combine saccule and utricle data for subsequent analysis to increase cell sample size and boost statistical power.

### Distinct regeneration responses of the zebrafish adult inner ear

Our scRNA-seq revealed regeneration responsive transcription in a cell type specific manner. Although Uniform Manifold Approximation and Projection’s (UMAPs) of individual cell types show that the cells in non-regenerating and regenerating tissues cluster together (Figure 3A), the regenerating inner ear exhibits an expansion of a class of cells that were intermediate between support cells and hair cells (“PC2” in Figure 3B) and each cell type possessed significant differences in the transcriptome compared to homeostatic controls. We tested whether the observed changes were caused by DT injection alone by performing pairwise comparisons between Day 4 WT and WT fish injected with DT. The homeostatic controls (Untreated DTR and Day 4 treated WT fish) cell populations were essentially identical and pooled for comparisons.

**Figure 3.**
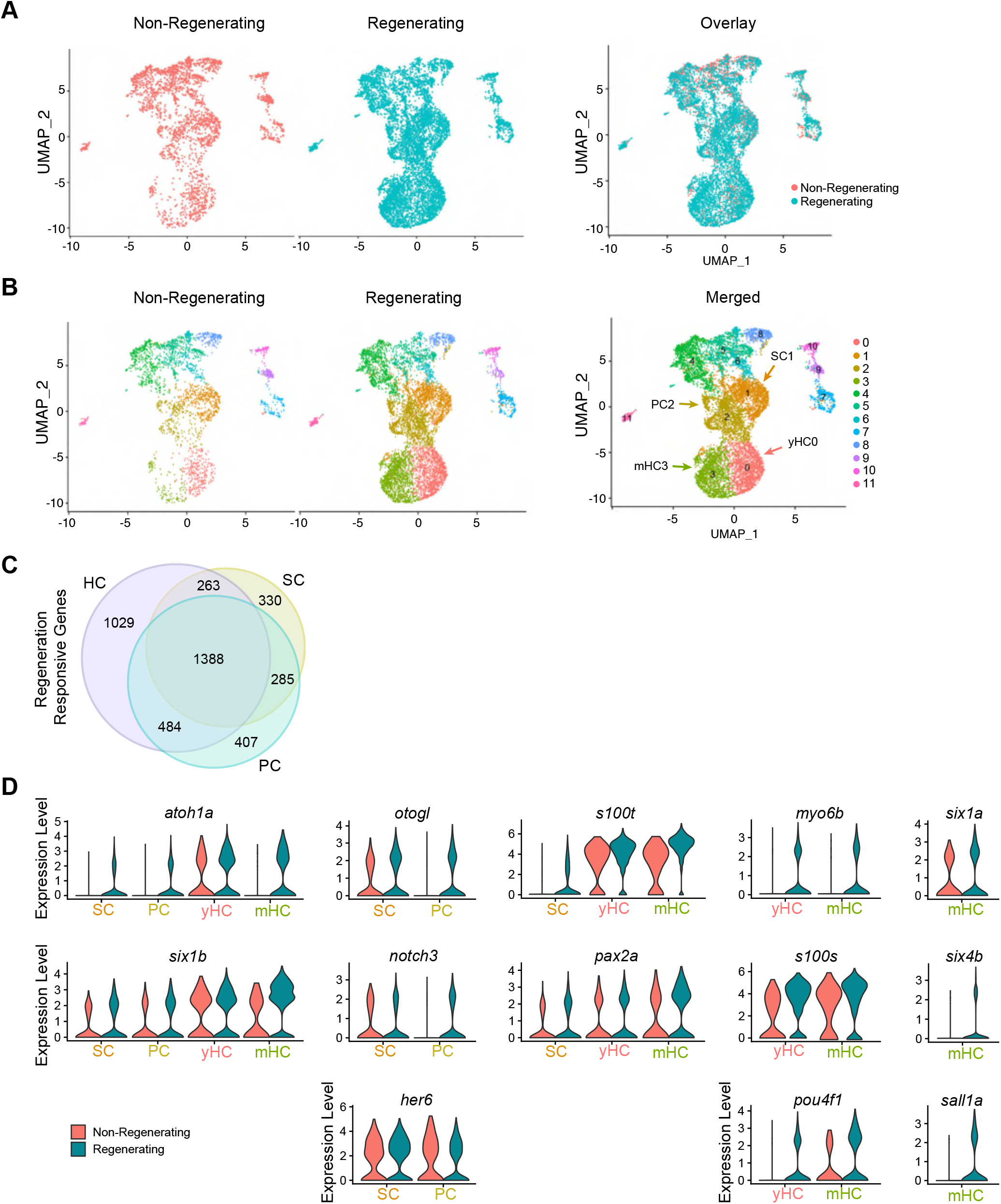
Gene expression during regeneration shows cell-type specific gene expression and expansion of the progenitor cell population. (A) UMAP plots of inner ear cells split between non-regenerating controls and regenerating sensory epithelia. Cells across conditions group together based on basal gene expression, allowing for a single joint clustering (B) to detect 12 cell populations. (C) Venn diagram showing the overlapping and unique differentially expressed genes for each cell type, hair cells “HC”, support cells “SC”, and progenitor cells “PC”. (D) Violin plots showing the distribution of gene expression of top genes identified with DE testing across cell types by comparison to non-regenerating cells.

We determined how many of those genes were differentially expressed as a consequence of regeneration. We identified 2,266 genes altered in expression between controls and regenerating support cells, 2,564 in progenitor cells, and 3,164 in hair cells (p-value < 0.1; FC ≥ 2). 1,388 differentially expressed genes were shared among all three cell types. 1,673 differentially expressed genes were shared between SC and PC, 1,872 between PC and HC, and 1,651 between SC and HC (Figure 3C Venn diagram, Table S7).

Global transcriptional changes in sensory epithelial cells involved in the hair cell lineage were observed following hair cell ablation. The genes *atoh1a* and *six1b* were upregulated in all regenerating sensory cell types – SC, PC, and HC types. Supporting and progenitor cells showed an upregulation of *her6* (mammalian *Hes1*), *notch3*, and *otogl*. Hair cell specific genes such as *s100t* and *pax2a* were upregulated in supporting cells and in young or mature hair cells. Young and maturing hair cells exhibited an upregulation of *myo6b*, *s100s*, and *pou4f1*. Finally, mature hair cells were marked by an upregulation in *six1a*, *six4b*, and *sal1a* (Figure 3D).

### Temporal gene regulation patterns in hair cell regeneration

We sought to visualize the continuous program of gene expression changes that occurs during hair cell differentiation. By reversed graph embedding, Monocle3 measures progress in “pseudotime” which is an associative measure of how much progress an individual cell has made through a process such as cellular regeneration or differentiation (Figure 4) (Trapnell et al., 2014). Monocle3 grouped cells involved in hair cell regeneration and differentiation into 8 clusters and cell types were assigned to each cluster based on its gene expression profile (Figure 4A).

**Figure 4.**
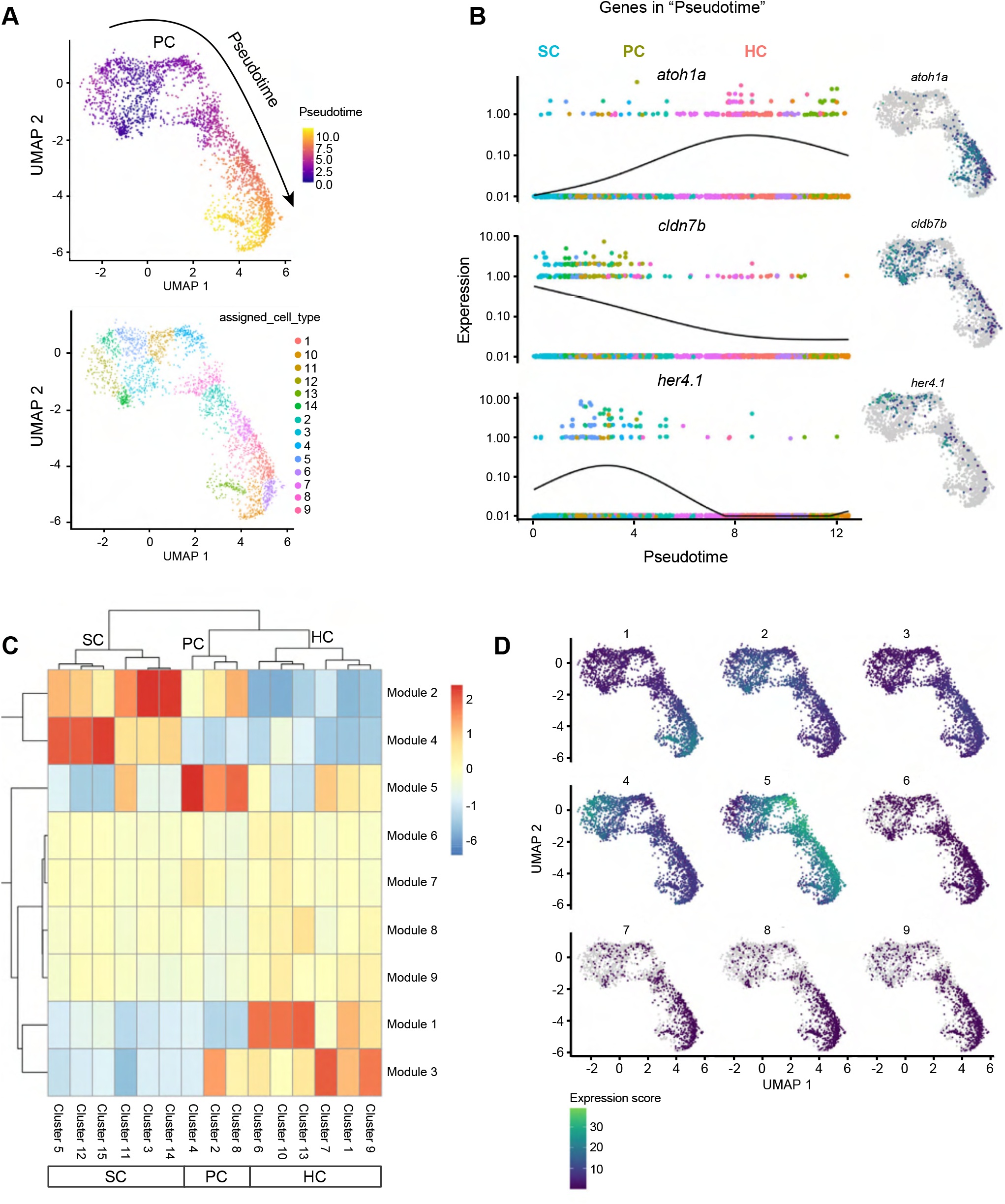
Cell lineages in the inner ear. (A) Top: Cells progressing through the regeneration program measured by Monocle3 in pseudotime. Arrow indicates the trajectory of the pseudotime pathway. Support cells are the root node of the trajectory graph, progenitor cells emerge after hair cell injury and are a clear intermediate to both support cells and hair cells. The point where support cells transition to progenitor cells is labeled “PC”. Bottom: Cluster breakdown of cells progressing in pseudotime. (B) Top differentially expressed gene for each cell type to show their dynamics as a function of pseudotime. (C) Gene modules co-regulated along the pseudotime during hair cell initiation, progression, and maturation. (D) Maps showing modules with specificity to specific cell clusters, while other modules are shared across multiple states of pseudotime. Top upregulated and downregulated GO terms are described in text for the module maps.

Using the trajectory graph, we ordered the cells according to their progress through the regeneration program (Figure 4A). The support cells possessed the earliest developmental stage assignment and were designated as the root of the trajectory. Pseudotime was calculated for all other cell types based on their distance from the root of the trajectory. Examples of genes that were expressed early (in support cells) and late (in mature hair cells) in pseudotime in the UMAP cluster are shown in Figure 4B. By building single-cell trajectories with scRNA-seq data, we observed switch-like changes in expression of key regulatory factors such as *cldn7b*, *her4.1* (*Hes5* in mammals), and *atoh1a*. The pseudotime ordering of cells showed that some genes act very early in support cells and then get shut off (like *cldn7b*) whereas others display dynamic temporal activity, turned on and then shut off in progenitor cells and/or newly specified hair cells.

We took all the genes that varied across the clusters (Figure 4A, bottom UMAP cluster) and grouped those with similar patterns of expression into gene modules (Figure 4C). We identified 9 co-expression modules in response to temporal changes in the hair cell regeneration program. These modules represented genes that share very similar expression patterns during hair cell regeneration and showed genes that were upregulated at various stages of the regeneration process (Figure 4D; Table S8). The identified modules of co-regulated genes were specific to certain clusters of cells. We explored the genes in each module and conducted GO enrichment analysis (Table S8).

Module 1 was specific to young hair cells, while module 3 was specific to mature hair cells. Genes grouped in module 1 were involved in *hair cell differentiation*, s*ensory perception of sound*, *microtubule-based movement*, *detection of mechanical stimulus involved in sensory perception*, *ATP biosynthetic process*, and *aerobic electron transport chain*. Module 3 genes were involved in *mRNA splicing*, *rRNA processing*, *proteasome-mediated ubiquitin-dependent protein catabolic process*, and *translation*. Module 5 was specific to progenitor cells and the genes were involved in *ribosome biosynthesis*.

Module 2 was specific to support cells and progenitor cells. Genes grouped in module 2 included both notch independent (*hes2.2*, *hes6*) and notch dependent (*her4.1*, *her4.2*, *her4.3*, *her4.4*) factors, *notch1a/b*, *sox10*, and *otogl.* The GO terms associated with Module 2 were *lateral line system development* and *cell junction organization/morphogenesis of epithelium.* Module 4 was highly specific to support cells. Genes grouped in module 4 include mammalian *HES1*-related genes, *her6* and *her9*, with the associated GO terms *negative regulation of transcription*, implying the increase of *her* repressors in progenitor cells during regeneration. We propose that modules 2 and 4 might be important for allowing the support cells and progenitor cells to make cell-state transitions (Soto et al., 2020).

### scATAC-seq reveals chromatin accessibility during regenerating of the zebrafish inner ear

We performed scATAC-seq to explore the chromatin accessibility landscape of the regenerating adult zebrafish inner ear. We obtained eleven scATAC-seq profiles using the 10X Chromium system and Cell Ranger ATAC 2.0.0 pipeline (Figure 5; Table S9). We assessed the quality of the dataset based on correlation with bulk ATAC-seq performed on similar tissues, the insert size distribution with peaks, and the relative enrichment of fragments around the TSS. We found that the aggregate of scATAC-seq profiles closely resemble bulk ATAC-seq samples, indicating that aggregate scATAC-seq captured the chromatin accessibility in a manner equivalent to bulk ATAC-seq assays (Figure 5A). A hallmark of high quality ATAC-seq libraries is a banded insert size distribution with peaks or genomic signals representing putative regulatory regions resulting from nucleosome protection, which was apparent even in individual cells (Figure 5B). This resulted in a dataset with 11 single-cell epigenomes (Figure 5B, UMAP plot).

**Figure 5.**
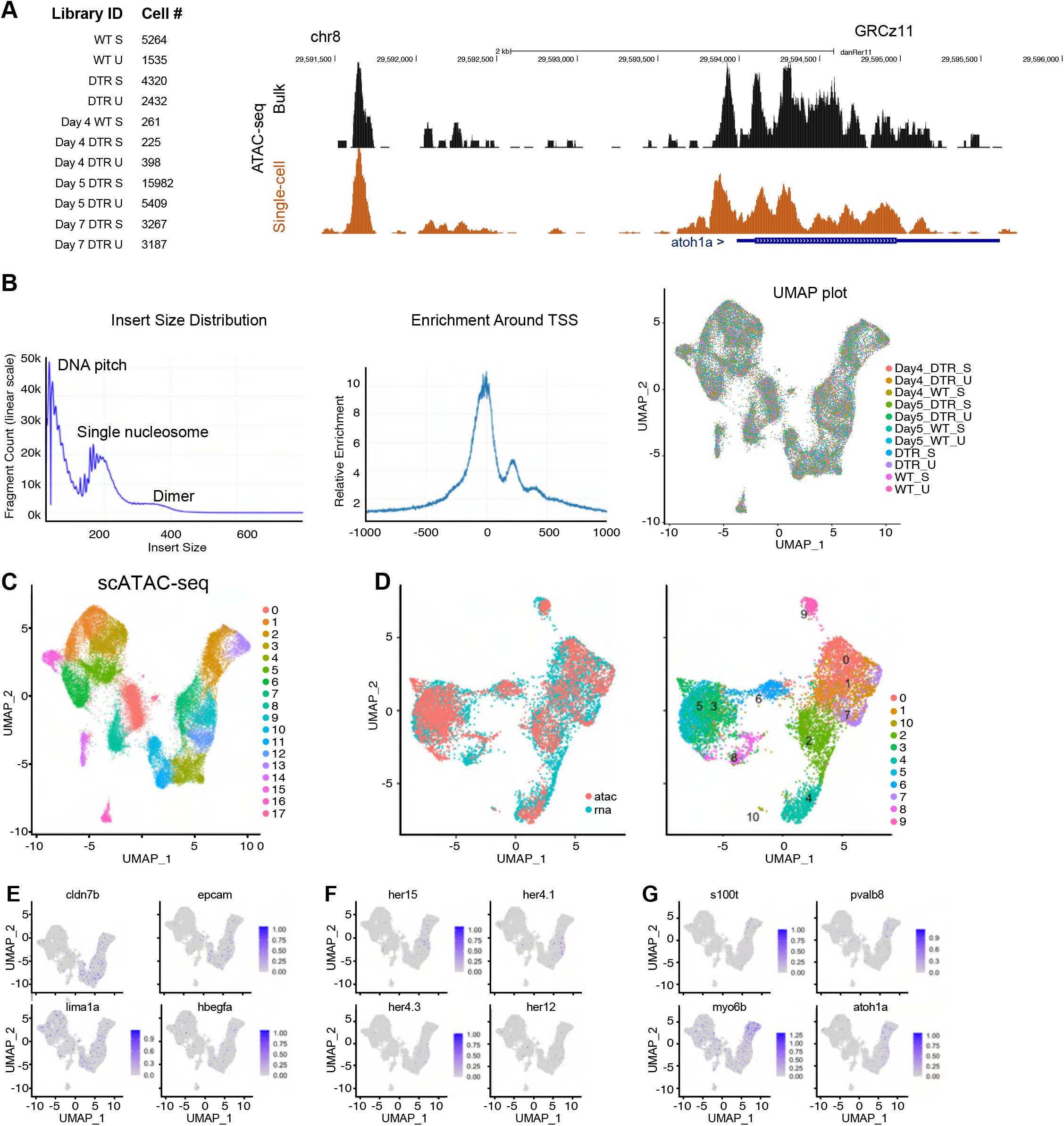
scATAC-seq of auditory and vestibular epithelia during regeneration. (A) Left: Identifiers and cell counts for samples collected for scATAC-seq. S = saccule, U = utricle, DTR = diphtheria toxin receptor, and Day = number of days after DT injection. Right: The aggregated scATAC-seq profiles resembled bulk ATAC-seq data from inner ear tissue. (B) Left: The library fragment size distribution displayed clear nucleosome banding patterns. Middle: Sequencing reads showed strong enrichment around transcriptional start sites (TSS). Right: We observed high sample agreement between all library samples, demonstrating the quality of the data was high. (C) Aggregate of all scATAC-seq samples including untreated samples. (D) Left: Integration shows concordance between chromatin accessibility (ATAC) around gene TSS and gene expression (RNA). Right: RNA cluster groups maintain cohesion in ATAC-seq data. ATAC-seq data still separates strongest marker genes correctly in (E) support cells, (F) progenitor cells, and (G) hair cells.

Using unsupervised clustering methods, single nuclei were clustered and cell types were identified based on quantitative accessibility and gene expression (Figure 5C; Table S10). Although the proportion of reads overlapping peaks was lower for some samples, the samples were well correlated. The total number of cells profiled ranged from 225 to 15,982 per sample. The average number of median unique fragments per cell in the library was 3,618. The average total mapped read pairs was 40,728,736. Overall, these results indicated that our scATAC-seq data were of high quality and suggested that the individual nuclei could reveal cell-specific patterns of the constituent cell types.

We next sought to annotate clusters. Many clusters were identifiable based on cluster specific promoter accessibility of the cell-type-specific genes. We validated the identity of our clusters by integrating scATAC-seq with scRNA-seq profiles (Figure 5D). We observed that our chromatin accessibility profiles across cell-type signature genes were enriched with our assigned scRNA-seq cluster identities (Figure 5D, right UMAP plot). For example, the promoters of *atoh1a* were specifically accessible in cluster 1 and likely belonged to the hair cell lineage (Figure 5G).

### Identification of regeneration-responsive elements (RREs)

We found dynamic changes in chromatin accessibility that emerged as a consequence of regeneration. We identified regions that presented higher accessibility during regeneration compared to controls and named these regions “regeneration-responsive elements” (RREs). To characterize RREs, we obtained single-cell ATAC-seq data from untreated inner ear saccules to represent the basal state of the tissues. We identified differentially accessible peaks between clusters of cells by performing a differential accessibility test (Seurat’s Signac). To identify RREs, we removed common peaks shared between untreated and treated samples leaving only the cell-type-specific emerging peaks.

Using this approach, we identified 12,369 RREs in supporting cells, 13,528 RREs in progenitor cells, and 12,597 RREs in hair cells (Figure 6A; Table S11). The chromatin accessibility of progenitor cells and hair cells showed a much higher degree of overlap than the support cells did with either progenitor cells or hair cells. To understand the functions of emerging peaks, we applied GREAT analysis to annotate peaks and predict functions of putative regulatory regions (Figure S3) (Hiller et al., 2013). GREAT analysis indicated that ∼19.5-25.0% of peaks were located in proximal regions of the transcriptional start, whereas ∼70-75% were located in distal regions (> +/- 5 kb) (19.46% proximal PC peaks, 24.98% proximal SC peaks, 21.12% proximal HC peaks), many of which were in the first introns of genes. GO functional annotation of emerging peaks identified that peaks from all three cell types were associated with *lateral line* and *inner ear development* or differentiation such as *mechanosensory lateral line system development* (p-value ≤ 6.2061e-6; SC peaks), *mechanoreceptor differentiation* (p-value ≤ 3.4662e-9, PC peaks), and *inner ear receptor cell differentiation* (p-value ≤ 1.4879e-8; HC peaks) (Figure S3 D, E and F). Our results demonstrate that the RRE accessible chromatin regions were highly associated with hair cell differentiation genes.

**Figure 6.**
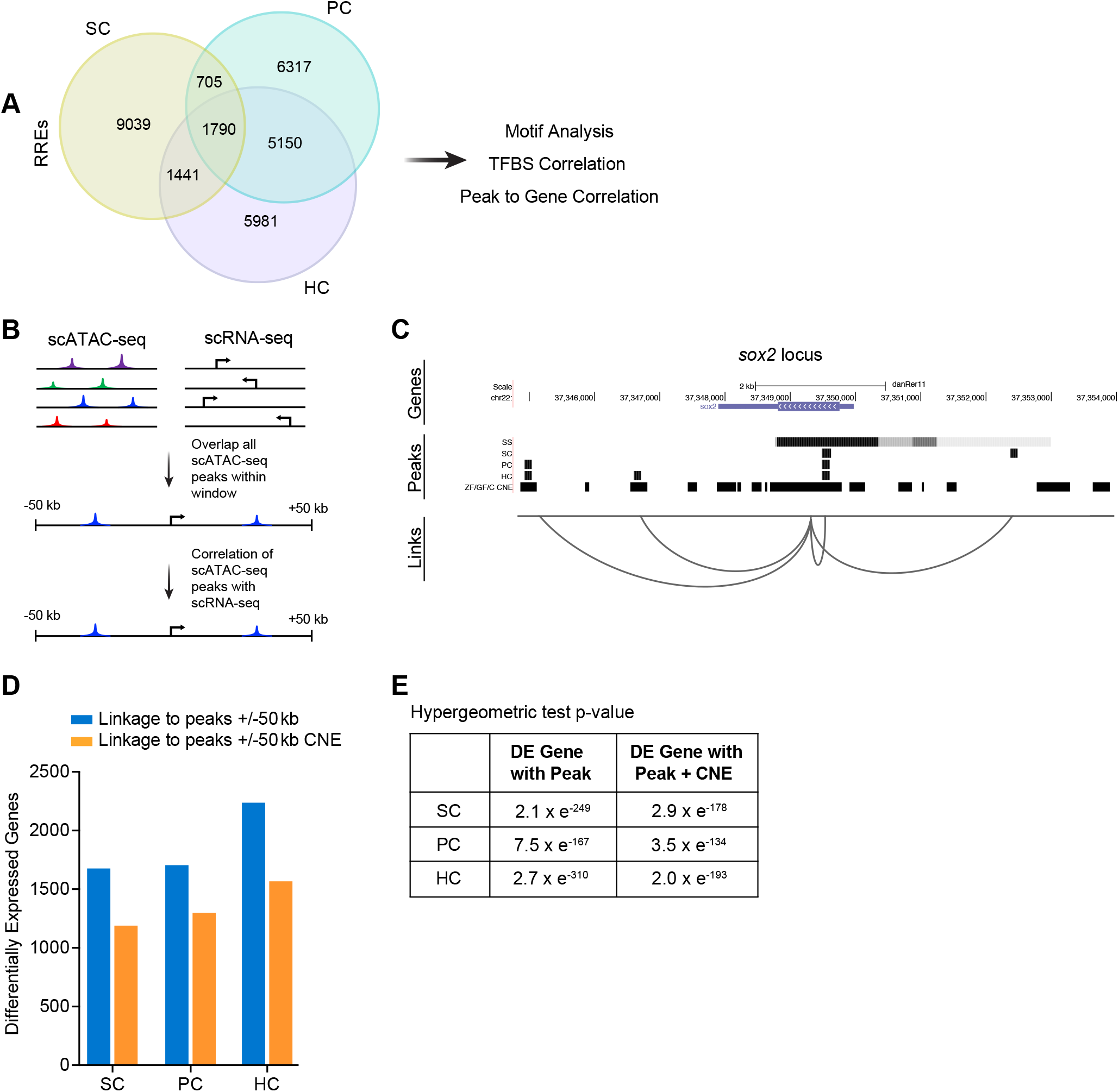
Correlation of scATAC-seq peaks to conserved regions and to differentially expressed genes. (A) Venn Diagram of emerging peaks identified in support cells, progenitor cells, and hair cells. (B) Schematic of the approach used to link scATAC-seq peaks in proximal and distal DNA elements to genes by correlating chromatin accessibility and differentially expressed genes (scRNA-seq). (C) Predicted links between peaks near the gene, *sox2*. For each cell type, a track is shown including peaks present in controls (ss: steady state) and CNEs. Peaks or genome coordinates are represented by black bars. (D) Bar graphs showing the distribution of RRE peaks intersected with differentially expressed genes and CNEs. (E) Intersections are statistically significant based on hypergeometric tests p-value <0.001.

To determine whether cell-specific open chromatin regions from scATAC-seq analysis correlated with cell-specific gene expression, we developed a computational strategy to infer enhancer-to-gene relationships. We investigated to what extent enhancers in a window surrounding the TSS of a gene (+/-50 kb from the TSS) predicted the expression of a gene (Figure 6B). There were 67.5% more genes associated with peaks when we use a 50 kb window in comparison to the traditional 2 kb window. To investigate how accessibility relates to gene expression, we identified RREs 50 kb upstream and downstream from TSS’s (Figure 6B). Figure 6C shows an example for the *sox2* locus. For hair cells 13,966 genes were associated with peaks, for support cells 13,709 genes were associated with peaks, and in progenitor cells 13,874 genes were associated with peaks (Table S12).

We next correlated RREs with changes in gene expression during regeneration. We reasoned that correlation between RREs with differentially transcribed genes during hair cell regeneration may be used to connect regulatory regions to their target genes. Indeed, we found a clear correlation between chromatin accessibility and nearby/distal gene expression (p-value < 0.05, hypergeometric Test) (Figure 6D and Figure 6E; Table S15). We identified, 1,546 differential genes in progenitor cells associated with at least one RRE peak (p-value < 7.5 x e^-167^, hypergeometric test) (Fury et al., 2006), 1,497 support cell genes were associated with at least one RRE peak (p-value < 2.0 x e^-249^, hypergeometric test), and 2,032 hair cell genes had at least one nearby RRE peak (p-value < 2.7 x e^-310^, hypergeometric test) (Table S14). For further analysis, we chose the set of RREs that were within 50 kb of a gene with differential expression (regardless of whether that gene was the closest gene to the RRE) assuming that this subset would be enriched for regulatory regions that are having a direct effect on gene expression levels.

### RREs are associated with conserved, non-coding elements

Conserved, non-coding elements (CNEs) are sequences outside of the coding regions that have a high degree of sequence conservation among multiple species. The “longer” a sequence is conserved across evolutionary time, the higher the probability it has a conserved functional purpose. Overlap of ATAC-seq peaks with CNEs implies functional significance (Wittkopp and Kalay, 2011). To further validate this data and likelihood of identified enhancers for functional roles *in vivo* and to also make use of evolutionary sequence conservation as a filter to find putative gene regulatory elements, we intersected identified RREs associated with differential gene expression with CNE data derived from comparisons between zebrafish and several carp species generated by Chen et al (Chen et al., 2019) (Figure 6D; Table S13). We found significant overlap between conserved noncoding blocks and RRE peaks associated with at least one differentially expressed gene identified in our scATAC-seq and scRNA-seq data (SC: p-value < 2.9 x e^-178^, PC: p-value < 3.5 x e^-134^, HC: p-value < 2.0 x e^-193^, hypergeometric test) (Figure 6E; Table S14 and S15). Together these data identified cis-regulatory elements strongly enriched for roles in transcriptional control during hair cell regeneration.

### Deep learning identifies key regulatory motifs and potential co-regulation cassettes

Machine learning can accurately identify enhancer sequences *de novo* from raw sequence data (Ahmad et al., 2014). Therefore, we constructed a prediction model to determine if enriched transcription factor binding sites (TFBSs) were found in the RREs. A deep learning model using a 59,785-parameter, 4-layer DL model consisting of convolutional, normalization, max pooling and dropout layers was trained by integrating features associated with enhancer activity such as TF binding from ChIP-seq, and these models were then applied to emerging peak sequences (Table 1). The model showed that Six and Sox motifs were significantly enriched in our datasets and the form of the enrichment was determined in a cell specific manner (Figure 7A). Furthermore, deep learning revealed that Sox factors were more strongly enriched in support cells (Figure 7B) and showed significant co-occurrence with Pdx1, Foxd3, Cebpa, Ebf1, and Rest.

**Figure 7.**
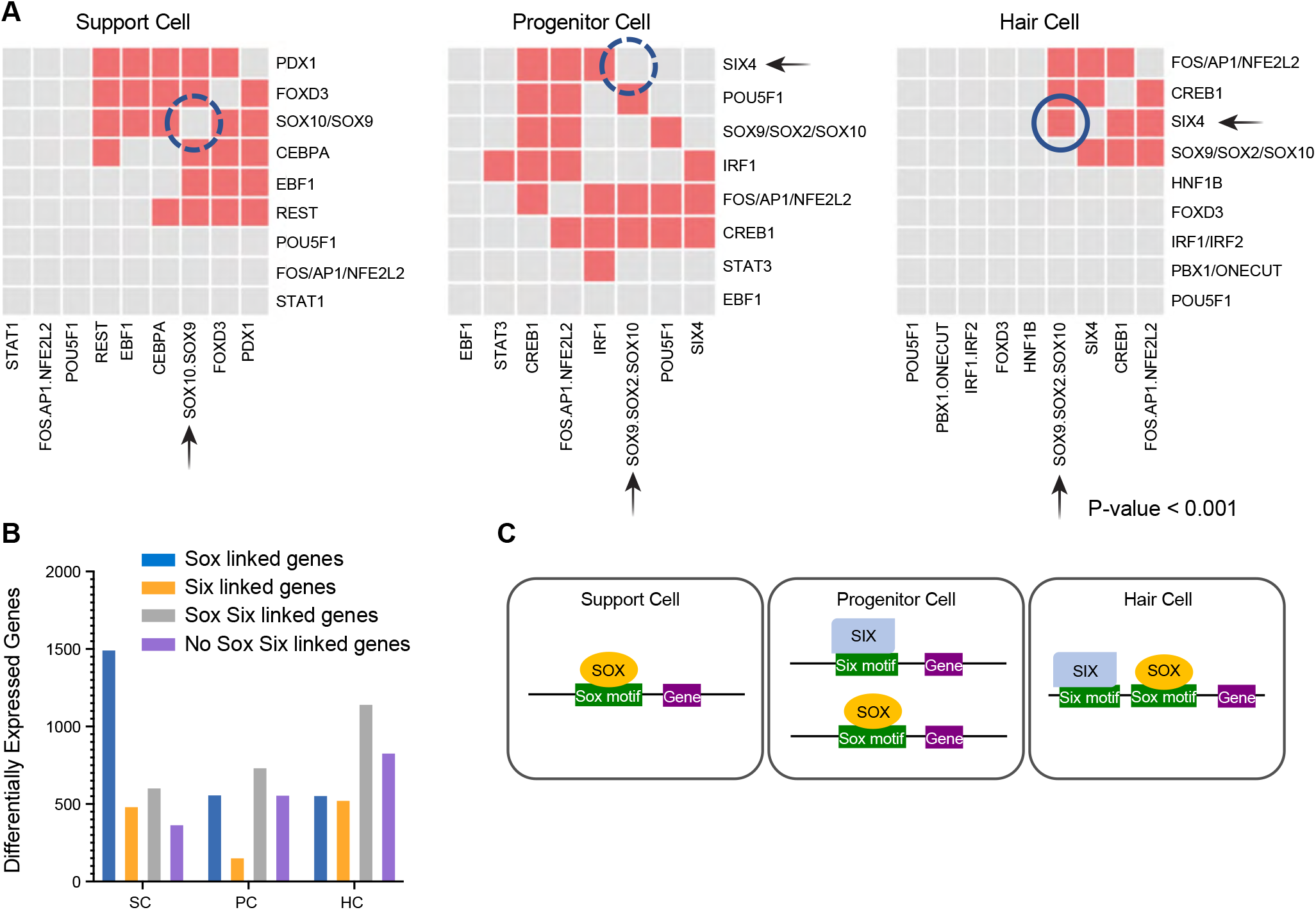
Combinatorial accessibility of Six and Sox transcription factors are defined by distinct cell identities. (A) Pair-wise TF co-association in support cells, progenitor cells, and hair cells. The colored boxes represent the extent of co-association between the TFs denoted in the rows and columns compared to chance. (B) Sox and/or Six TF motif containing enhancers associated with differentially expressed genes in support cells (SC), progenitor cells (PC), or hair cells (HC). Categories of TF containing motifs include Sox containing, Six containing, Sox and Six containing, or neither. Sox containing peaks connected to differentially expressed genes dominates the SC, while the largest category of linked peak in HC contained both Sox and Six binding sites. (C) Model of Six and Sox TFs with cooperative roles in mediating activity of individual regulatory elements such that the SIX and SOX are recruited to specific regulatory elements and function as either stem cell promoting or hair cell promoting factors.

**Table 1.** Convolutional layers and parameters used in supervised deep learning models.

| Structure of CNNs for Deep Learning model |  |  |  |
| --- | --- | --- | --- |
| Convolution1D(64, 9) | activation='relu' | kernel_regularizer=L1L2(l1=1e-4, l2=1e-3) | kernel_constraint=max_norm(1) |
|  | BatchNormalization() | MaxPooling1D(9, 3) | Dropout(0.2) |
| Convolution1D(32, 4) | activation='relu' | kernel_regularizer=L1L2(l1=1e-4, l2=1e-3) | kernel_constraint=max_norm(1) |
|  | MaxPooling1D(4, 2) | Dropout(0.2) |  |
| Convolution1D(32, 4) | activation='relu' | kernel_regularizer=L1L2(l1=1e-4, l2=1e-3) |  |
|  | MaxPooling1D(4, 3) | Dropout(0.2) |  |
| Convolution1D(32, 4) | activation='relu' | kernel_regularizer=L1L2(l1=1e-4, l2=1e-3) | kernel_constraint=max_norm(1) |
|  | MaxPooling1D(4, 2) | Dropout(0.5) |  |
| Flatten() | Dense(180, activation='relu') | Dense(1, activation='sigmoid') |  |
| loss='binary_crossentropy', optimizer='ADAM', metrics=['accuracy'] |  |  |  |

In progenitor cells, Sox and Six sites were more evenly enriched and both often co-occurred with Creb1 or Fos/Ap1 but not with each other. In hair cells, Sox sites showed significant co-occupancy with Six4 as well as with Creb1, and Fos/Ap1 (correlation of TFBSs p-value < 0.001) (Figure 7A). We parsed the RREs into four categories, containing only Sox motifs, containing only Six motifs, containing both Sox and Six motifs, or containing neither Sox nor Six motifs (Figure 7B). We then looked at how many of the peaks in each category were associated with differentially expressed genes in SC, PC, or HC (Table S16). We detected a clear pattern where in SC the strongest association with differentially expressed genes were Sox containing enhancers (1,498 genes) while Six, Sox/Six and Neither had 487, 608, and 370 associated genes respectively. It is notable that 71% of all enhancers linked to differentially expressed genes in the SC had Sox motifs. In PC, despite having more total peaks than SC, had 944 fewer gene associations total with no clear pattern of preference across the four categories except for Six-containing peaks being relatively poorly represented. This pattern may be consistent with a cell type that is in transition. HC had the most total differentially expressed genes associated with enhancer elements total, with the largest category being peaks containing both Sox and Six motifs (37%). The second largest category for HC had neither Sox nor Six motifs. In SC, the top GO enrichment terms for Sox-linked genes were: “negative regulation of protein processing,” “negative regulation of protein maturation,” “hemidesmosome assembly,” and “regulation of epidermis development” consistent with epithelial cells that would be undergoing a state transition. Highest GO enrichment for the HC Six/Sox-linked genes were: “peripheral nervous system axonogenesis,” “peripheral nervous system development,” and “peripheral nervous system differentiation” consistent with driving hair cell fates.

Based on the scRNA-seq data, we determined which Sox and Six factors were expressed in each cell type to correlate accessible binding sites to available transcription factors (Figure 8). The Sox genes *sox2*, *sox4b*, *sox10*, and *sox21a*, all showed expression changes, as did *six1a*, *six1b*, *six4a*, and *six4b*. The changes in *sox2* and *sox21a* were particularly restricted to the progenitor cells suggesting they are key drivers of the state change, while both *six1* genes (but particularly *six1b* in both regenerating and non-regenerating sets) were very strongly expressed in hair cells.

**Figure 8.**
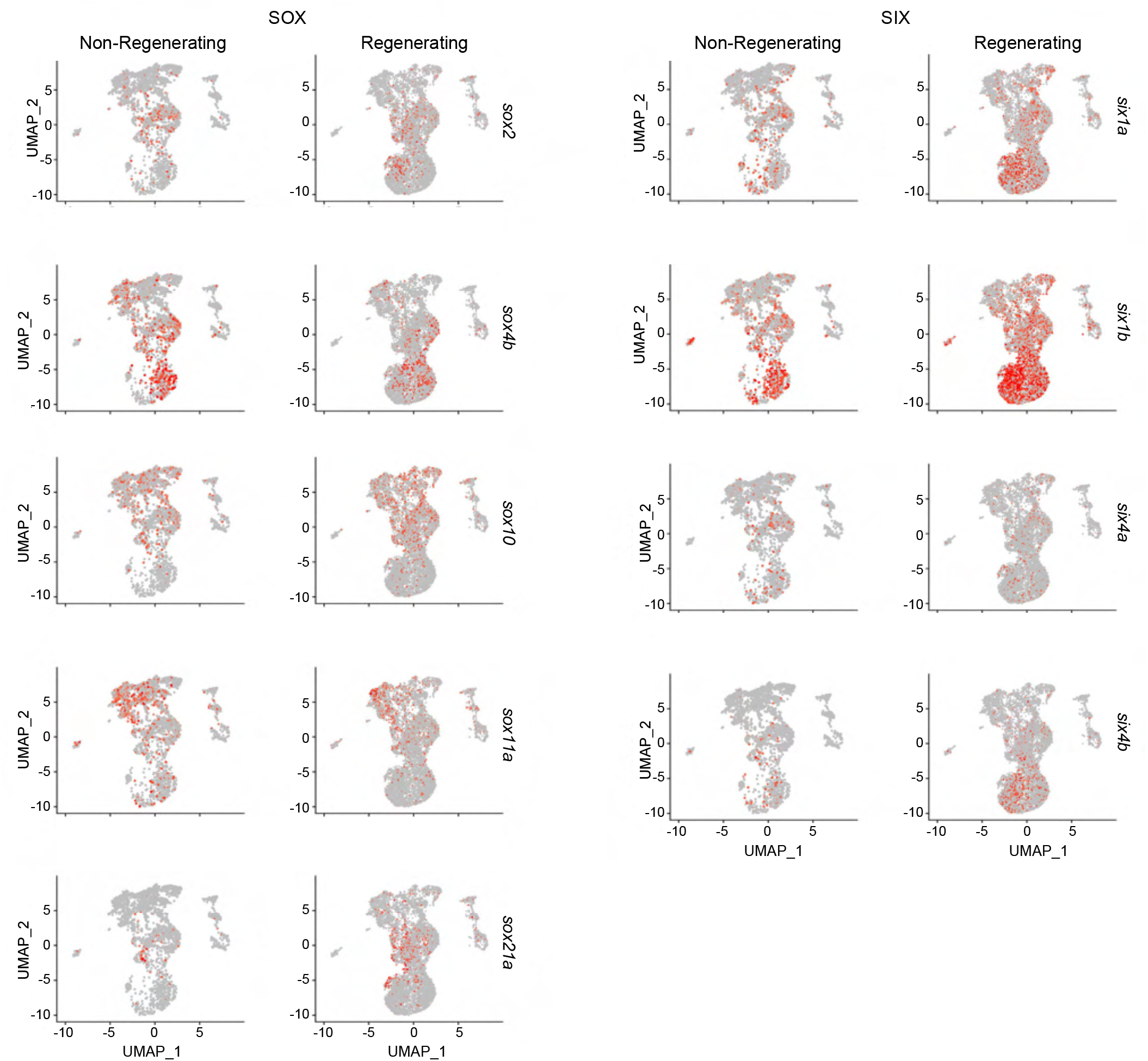
Sox and Six TF enrichment corresponds to changes in gene expression during hair cell regeneration. Comparisons between non-regenerating and regenerating inner ear tissues for the identified Sox and Six transcription factors identified as differentially regulated. The enrichment of Sox and Six motifs in RREs identified by machine learning correlated to expression of Sox and Six transcription factors during regeneration.

Six family members have a demonstrated role in mammalian hair cell differentiation (Zhang et al., 2017) and that knockout of *Six1* results in defective inner ear development (Zheng et al., 2003). In mice, *Sox2* has known roles in sensorineural development and is expressed in hair cell progenitors and support cells until it is downregulated after differentiation (Kempfle et al., 2016) and Sox genes in general are associated with pluripotent cells. The observed pattern of motif enrichment suggests a significant shift from Sox driven gene expression in SC to Sox/Six co-regulated expression in HC. The relatively smaller number of differentially expressed genes linked to the emerging peaks in the PC is consistent with them being a transient state between the SC and HC, where the chromatin is opening to allow for gene expression, but actual gene activation has not occurred yet. Once gene expression is initiated, the cells shift to a HC classification.

### Deletion of a predicted regulatory element of sox2 results in a dominant loss of hair cell regeneration

Our integrated single cell epigenomic and transcriptomic analysis predicted that *sox2* is differentially regulated in a switchlike pattern such that *sox2* expression turns on in progenitor cells and then shut off in newly differentiating hair cells (Figure S4). Based on the known role of *sox2* in stem cell pluripotency in general and its role in inner ear development specifically, we were interested in identifying if there were regulatory elements for *sox2* that were specifically activated during hearing regeneration and if *sox2* was a key driver of regeneration. We found the *sox2* locus acquired cell specific dynamic changes in accessibility during hair cell regeneration with ATAC accessibility of specific regions correlating to upregulation in gene expression (Figure 6C). A specific element 1995 bp upstream the TSS of *sox2* emerged in support cells and progenitor cells only during regeneration (but was absent in the hair cells), suggesting that this might represent an enhancer of *sox2* with important roles in regulating regeneration (Figure S5).

We used Seurat Signac to narrow down the peak to a 2,115 bp region. The peak overlapped with a CNE identified from zebrafish/goldfish/carp alignments (Chen et al., 2019) as well as two shorter “ultraconserved” regions present in all vertebrates (Siepel et al., 2005)(Figure S5). In addition, there were a number of TFBS predictions according to the JASPAR database (Khan et al., 2018) including sites for *sox2*, *pou4f3*, and *stat3* all of which are associated with inner ear development and/or regeneration.

To functionally validate if this region regulates *sox2* during regeneration, we utilized CRISPR/Cas9 to introduce deletions of the 2 kb *sox2* upstream enhancer. We designed and synthesized six guide RNAs to target the loci (three to a side) and co-injected them with spCas9 protein into single cell-stage embryos (Figure 9A). We identified germline transmitted deletions and generated heterozygous and homozygous mutant deletion lines. Experiments were performed on stable lines of heterozygous enhancer deletions or homozygous deletions, denoted here as *sox2^hg138^*.

**Figure 9.**
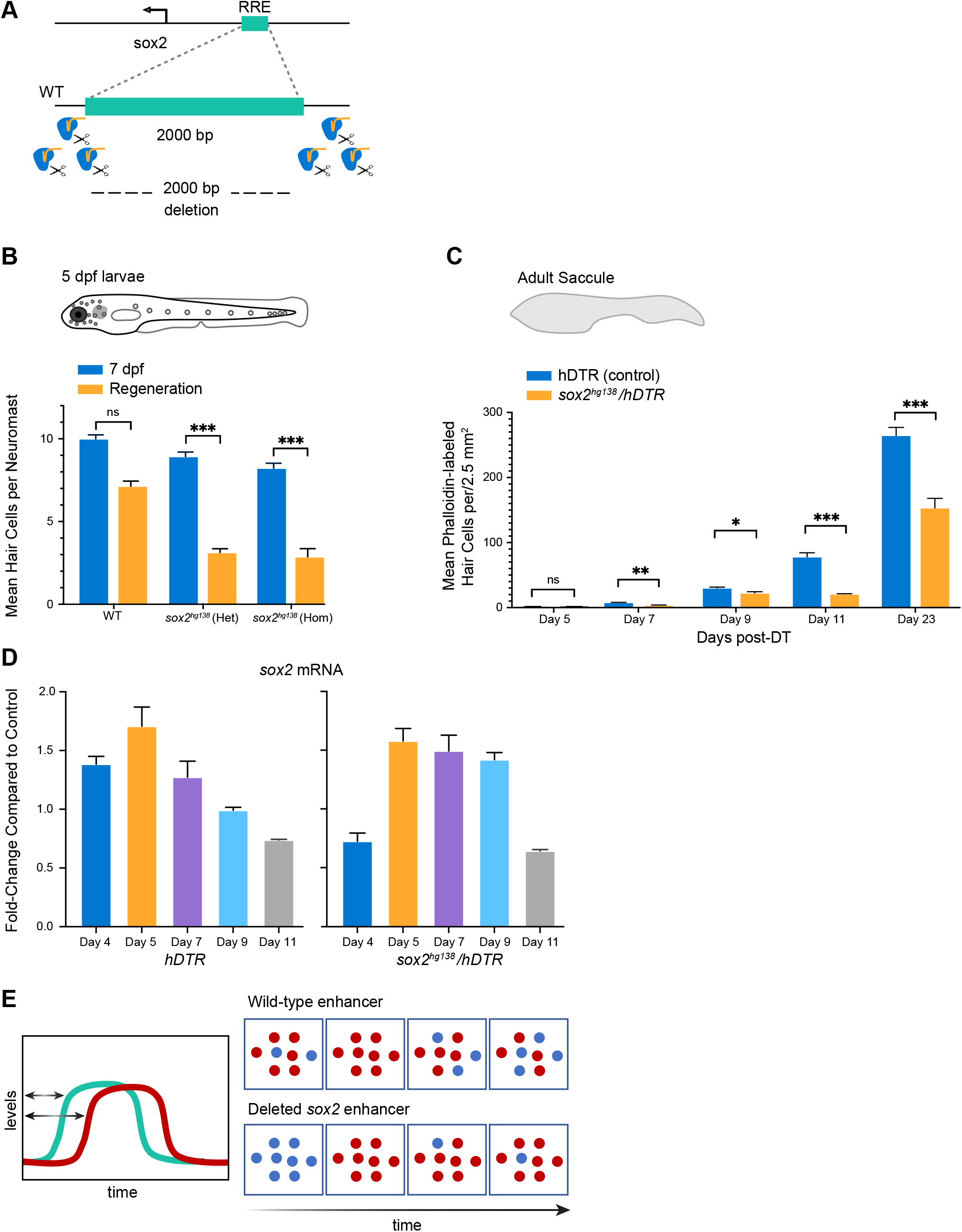
The –1995 bp *sox2* enhancer element is required for hair cell regeneration. **(A)** Generation of enhancer deletion mutants. Schematic diagram showing the deletion of the targeted enhancer using CRISPR/Cas9 editing. (B) Lateral line hair cell regeneration is strongly inhibited 2 days after ablation with CuSO_4_. (C) Adult hair cell regeneration is significantly inhibited up to 23 days after hair cell ablation. (D) Quantitative PCR measuring *sox2* mRNA levels in regenerating sensory epithelia and sensory epithelia from enhancer deletion mutants shows that the increase in *sox2* expression is delayed by 24 hours then remains elevated for 48 hours (day 7 and day 9). ns = not significant, * = p-value ≤ 0.05, ** = p-value ≤ 0.01, *** = p-value ≤ 0.001.

Enhancer deletion mutants had no overt morphological phenotypes in early larvae, and both heterozygous and homozygous deletions survived to adulthood. Homozygous and heterozygous fish were viable as adults and the overall morphologies of inner ears, sensory epithelia, lateral line neuromasts, and even swimming behaviors appeared normal. To investigate the regenerative abilities of animals heterozygous and homozygous for the upstream deletion, we performed larval lateral line regeneration assays on the enhancer deletion mutants using CuSO_4_ ablation. We found that both homozygous and heterozygous deletions had no effect on larval hair cell development but altered hair cell regeneration with similar severity regardless of whether the mutation was in one copy or two (Figure 9B) (two-way ANOVA p-value < 0.0001). Therefore, the enhancer region was a haploinsufficient regulator of neuromast hair cell regeneration.

To examine the role for the *sox2* upstream enhancer deletion in adult zebrafish, we again employed the Tg(*myo6b*:DTR) transgenic zebrafish (Jimenez et al., 2021) (Figure 9C). In our experiments, heterozygous *sox2^hg138^*/Tg(*myo6b*:hDTR) and controls (Tg(*myo6b*:hDTR) and WT) zebrafish received a single 0.05 ng/µL injection of DT, and sensory epithelia were examined after 7, 9, 11, and 23-days of recovery following injection. Fixed whole mount saccules were labeled for phalloidin and imaged using confocal microscopy. In untreated heterozygous *sox2^hg138^*/Tg(*myo6b*:hDTR), saccular hair cells were present in normal numbers (Figure S6). In DT injected heterozygous *sox2^hg138^*/Tg(*myo6b*:hDTR) zebrafish, there was a major reduction in hair cell regeneration after DT treatment on days 7, 9, 11, and 23 post DT in comparison to normally regenerating sensory epithelia (two-way ANOVA p-value < 0.0001). Our data reveal that the –1995 bp upstream enhancer of *sox2* is required specifically for hair cell regeneration but not normal hair cell development in zebrafish and heterozygous deletions possessing a phenotype as severe as homozygous deletions. We did not notice any regeneration deficits after tailfin amputation in the *sox2^hg138^*/Tg(*myo6b*:hDTR) fish, suggesting the regeneration phenotype for this deletion was restricted to hair cell regeneration, or perhaps to neuronal regeneration in general.

### The -1955 bp sox2 enhancer regulates the timing of sox2 expression during regeneration

We sought to determine if *sox2* expression levels were altered in both the heterozygous and homozygous *sox2* enhancer deletion fish in comparison to controls during regeneration. *sox2* mRNA levels were measured in sensory epithelia by quantitative RT-PCR. In non-regenerating sensory epithelia of the enhancer deletion mutants, *sox2* RNA levels were not significantly altered in comparison to control sensory epithelia. In adult zebrafish without the enhancer deletion undergoing hair cell regeneration, *sox2* expression was elevated on days 4, 5, and 7 post-DT. This was consistent with our single cell transcriptomics assays on regenerating inner ear tissues collected at all timepoints (Days 4,5,7) (Figure 8). By day 9 post-DT, *sox2* expression levels were near control levels and were further reduced on day 11 post-DT (Figure 9D).

In both heterozygous and homozygous enhancer deletion mutants, we found that *sox2* expression levels were significantly reduced on day 4 in comparison to normal inner ear sensory epithelia undergoing regeneration (p-value ≤ 0.0039). By day 5-7 post DT, *sox2* expression levels rose to levels comparable to normally regenerating inner ears. At day 9 post-DT, levels of *sox2* remain elevated in enhancer deletion mutants (p-value < 0.0021) in comparison to wild-type, but by day 11 post-DT, levels of *sox2* did drop back to levels comparable to wild-type. Our data suggest that the upstream enhancer of *sox2* may be specifically involved in regulating the timing of *sox2* expression but not completely essential for triggering activation (Figure 9E). It is interesting to note that despite *sox2* levels apparently only being shifted by 24 hours in the mutant compared to the wild-type, the regeneration of hair cells appeared to have been affected out to at least 23 days post ablation and potentially permanently.

## DISCUSSION

Regeneration inducible or injury dependent gene expression may be controlled by specific enhancer regulatory elements (Begeman et al., 2020; Goldman et al., 2017; Harris et al., 2016; Kang et al., 2016; Rodriguez and Kang, 2020; Soukup et al., 2019; Suzuki and Ochi, 2020; Thompson et al., 2020; Vizcaya-Molina et al., 2018; Wang et al., 2020). Tissue regeneration enhancer elements, or TREEs, have been identified by approaches such as H3.3 profiling and epigenetic profiling. While validation of identified regulatory elements revealed that the putative enhancers can direct expression of minimal promoters or reporters, deletion of multiple putative enhancers from the genome previously resulted in no detectable effects on regeneration (Goldman et al., 2017; Kang et al., 2016).

Here we used a single-cell profiling approach to identify regeneration responsive enhancers that emerge as a consequence of hair cell regeneration in the adult zebrafish inner ear sensory epithelia and correlated genome accessibility with changes in gene expression. We integrated cell-type, enhancer activation, and TF expression to understand the regulatory logic behind zebrafish hair cell regeneration. We demonstrated that regulatory elements were differentially active in each cell type during hair cell regeneration and represented specific cell phases, indicating that regulatory plasticity may be a key facilitator triggering the adaptive reprogramming of supporting cells to generate new hair cells. Our single-cell ATAC-seq data revealed regions within the zebrafish genome with chromatin accessibility signatures that were key regulators of gene expression specifically during hair cell regeneration.

### Origin of hair cells during zebrafish inner ear hair cell regeneration

Our regeneration assay causes selective hair cell ablation, leaving the population of differentiated Sox2 positive supporting cells undamaged (Figure 1). Similar to what is observed in the regenerating chick cochlea following acoustic trauma (Girod et al., 1989) and consistent with clonal analysis of HC origins in the chicken hearing organ the basilar papilla and lateral line organs in zebrafish (Fekete et al., 1998; Ma et al., 2008), our single-cell trajectory analysis shows that supporting cells (SCs) contribute new hair cells in the zebrafish inner ear by reverting to a less differentiated state we have labeled as progenitor cells. The progenitor cells represent a special transition state cell type that can reenter mitosis when hair cells are depleted and either choose to become a new hair cell or return to support cell identity.

There is some evidence for the presence of inner ear cell types with the capacity for self-renewal and hair cell-like formation in the mammalian vestibular organ of mice (Li et al., 2003). However, the adult cochlea completely lacks the capacity to regenerate due to the loss of a stem cell population as the mammal ages (Oshima et al., 2007). The continued effort to reprogram differentiated mammalian inner ear cells may require comparing how support cells propagate new hair cells in response to damage in regenerating model systems to that of the non-regenerative mammalian counterparts. Identifying enhancers that are key regulators of regeneration are an essential step in identifying the differences in regenerative capacity.

### Six and Sox TFs appear to act in a combinatorial fashion to drive cellular identity

The *Sox* and *Six* genes are known to be among highly expressed TFs important for hair cell development and differentiation (Iyer and Groves, 2021). By correlating cell-type, enhancer activation, and TF expression in zebrafish adult inner ears during regeneration, our analysis revealed that hair cell regeneration is orchestrated by specific combinations of TFs (Figure 7A). The pattern of motif enrichment and co-occupancy suggests that Six and Sox TFs may have cooperative roles in mediating activity of individual regulatory elements such that Six and Sox are recruited to the specific sequences and may function as either stem cell-promoting or hair cell-promoting factors (Figure 7C). Our findings are consistent with mouse prosensory data: In mice, prosensory cells (Sox2-EGFP+) from the embryonic cochlear duct are enriched for motifs corresponding to Six, Sox, Gata, Ebf and Tead families as well as motifs for Grhl2, Lef1, Irf4 and Rest (Wilkerson et al., 2019). We identified 30,423 open chromatin regions enriched for many of the same transcription factors as in the mouse prosensory cells. Motif enrichment and co-occupancy that was evident in a cell type specific fashion in our data provides evidence for shifting roles of Six and Sox at different stages of hair cell regeneration and implicates several other transcription factor families to have a role in hair cell development and regeneration.

Our scRNA-seq trajectory analysis suggested that most Sox TFs act early in support cells and then get shut off while others such as *sox2*, *sox13*, and *sox21a* are observed to toggle on and off again (Figure S4). As for the Six TFs, single-cell trajectory analysis showed that most turn on late when hair cells are differentiating, particularly *six1b*, suggesting that the Six TFs may function as the molecular switch to drive hair cell progenitors towards a differentiated state. Our findings also suggest that Sox and Six may have other interacting partners that are recruited to drive different lineages or cell fate decisions (Figure 7A).

### Enhancer regulatory elements with roles in regeneration

Regeneration competent animals are likely to possess conserved genomic regions necessary to activate injury and regeneration responses (Wang et al., 2020), we made use of previously generated CNE data across zebrafish and several carp species to find that there was a large degree of overlap between conserved noncoding blocks with RRE peaks associated with differentially expressed genes. We identified thousands of RREs that were conserved for more that 60 million years of divergent evolution.

Previously generated enhancer deletions in zebrafish have had no detectable effects on regeneration responsive gene expression nor resulted in a regeneration defect (Thompson et al., 2020). The *leptin-b* enhancer in the zebrafish (Kang et al., 2016) and the WNT gene cluster BRV118 enhancer in the fruit fly (Harris et al., 2016) modulate gene expression after injury. However, ablation of both have been shown to be dispensable for regeneration. In this study, we characterized regions that emerge in response to hair cell regeneration and identified an upstream enhancer of *sox2*. Deletion of this region in single or double knockouts resulted in abnormal regeneration and mis-regulated expression of *sox2,* suggesting that the upstream enhancer is genetically required for normal hair cell regeneration and controls the timing of *sox2* expression. These observations were true for both the inner ear sensory epithelium and the lateral line. Our results demonstrate that not only are changes in sox2 expression an essential component of hearing regeneration, initiation of *sox2* expression alone was insufficient for regeneration and that the timing of initiation and the duration of expression for the *sox2* transcription factor were also critical factors for successful hearing regeneration.

The success of regeneration in zebrafish is arguably due to regeneration responsive regulatory regions as well as more generalized injury responses upon hair cell damage. Addressing if inner ear regeneration enhancer elements are different among species and that the capacity to regenerate a given tissue might be impacted by these sequence differences will provide insight on how to proceed in “reactivating” the regeneration response in mammals.

## Supporting information

All supplemental tables

## ACKNOWLEDGEMENTS

This research was supported by the Intramural Research Program of the National Human Genome Research Institute (ZIAHG200386-06). We thank Dr. Tannia Clark and Charles River for zebrafish care; Suiyuan Zhang and the Bioinformatics and Scientific Programming Core; Stephen Wincovitch and the NHGRI Cytogenetic and Microscopy Core Facility; Martha Kirby, Stacie Anderson and NHGRI Flow Cytometry Core Facility; Di Huang for her valuable suggestions on computer programming and the computational resources of the NIH HPC Biowulf cluster (http://hpc.nih.gov); and the members of the Burgess laboratory for helpful discussion. All animal experiments were approved by the National Human Genome Research Institute’s Animal Care and Use Committee (protocol #G-01-3).

## AUTHOR CONTRIBUTIONS

EJ and SMB designed the experiments and wrote the manuscript. EJ, CS, and WS performed experiments. Single-cell genomics sample processing and sequencing were performed by WW and AGE. Bioinformatics analysis support were provided by ZC and DG. Animal support during DT studies were provided by SF. IO and SMB provided resource support.

## MATERIALS AND METHODS

### Experimental Animals

TAB5 (wild-type, WT), heterozygous Tg(*myo6b*:hDTR), and *sox2^hg138^* zebrafish used in this study were housed and raised on a recirculating aquaria system, using methods and parameters previously described (Frederickson et al., 2021). Fish were randomly selected and represented roughly equal numbers of males and females for single-cell genomics experiments. All experiments were approved by the Institutional Animal Care and Use Committee for the National Human Genome Research Institute under Animal Study Protocol: G-01-3.

### Adult Zebrafish Diphtheria Toxin Administration

Diphtheria toxin was purchased from Sigma-Aldrich (D0564) and dissolved in 1XPBS. 6 to 10 month-old wild-type (TAB5) and transgenic adult zebrafish of mixed sex were given one intraperitoneal (IP) injection with diphtheria toxin using a 10 µL NanoFil^TM^ microsyringe (NANOFIL) with a 35G beveled needle (NF35BV-2). Total protein injected into each fish was 0.05 ng in a total volume of 1 µL. Fish were fasted for 24 hours prior to IP injection. Buffered tricaine (<0.04 g/L) diluted in aquaria water was used to immobilize the fish, they were then placed (inverted) into a cut sponge and given the IP injection into the abdominal cavity, posterior to the pelvic girdle. Immediately after injection, fish were recovered in fresh system water and maintained off system for up to 23 days at a maximum density of 5 fish/L. Fish were fed Gemma 300 at approximately 1.5% body weight daily. Water quality (pH, ammonia, nitrite, nitrate, temperature) was monitored twice daily. At least 50% of the water was changed once daily. Health monitoring was performed twice daily and any fish that appeared to be in pain or distress were euthanized. Tissues were harvested on days 4, 5, and 7 for profiling experiments and over a broader range for other experiments.

### 10X Genomics scRNA-seq and scATAC-seq

Saccule and utricle were harvested from at least 35 adult zebrafish on days 4, 5, and 7 post-DT. Dissected sensory epithelia were dissociated for single-cell experiments using a previously described protocol with 1% BSA used in place of FBS (Bresciani et al., 2018). Cell number and viability was determined using a Luna Cell Counter.

For scRNA-seq, gel Bead-in-Emulsions were prepared by loading up to 7,000 cells per sample in 46.6 µL DMEM + 1% BSA onto a Chromium Chip B (10x Genomics 1000073) and run using the Chromium Controller (10x Genomics). cDNA libraries were generated with Chromium Single Cell 3’ GEM, Library and Gel Bead Kit V3 (10x Genomics). Libraries were sequenced using the NextSeq 500/550 High Output Kit v2.5 (Illumina) on an Illumina NextSeq 550. Reads were aligned to the zebrafish genome and feature-barcode matrices were generated using Cell Ranger 6.0.0 (10x Genomics). All scRNA data was deposited to GEO under accession GSE192947. UMAPs and differential gene expression lists were generated using Seurat. Downstream bioinformatic analysis was performed in R using the package Seurat (v4.0) (Butler et al., 2018; Stuart et al., 2019). UMAP dimensional reduction was performed on scaled data using Seurat function RunUMAP. Cell clusters were identified using Seurat function FindClusters. Markers to define clusters via differential expression were identified using the Seurat function FindAllMarkers.

To construct single cell trajectories, Monocle3 was used to cluster cells and reduce the dimensionality of the data using UMAP (Becht et al., 2018; Cao et al., 2019; Qiu et al., 2017; Trapnell et al., 2014). After identifying the trajectory graph of clustered cells, cells were ordered according to their progress through the developmental program and measured in pseudotime. We identified sets of genes that varied across clusters and Monocle3 grouped them into modules according to similar patterns of expression using the gene_module_df function.

Following integration between saccule and utricle experiments, conserved cell type markers were identified using the FindConservedMarkers() function. To identify differentially expressed genes across saccule and utricle for cells of the same type, we used the FindMarkers() function. Differential genes used for downstream analysis were required to have a p-value < 0.1 and an average log fold change of 0.25 in at least one dataset.

For scATAC-seq, nuclei were prepared from cell suspensions according to 10X Genomics Chromium sample preparation guidelines. The resulting nuclei concentration was determined using a Luna Cell Counter. Nuclei were adjusted to the desired capture number concentration based on the number of available nuclei. Up to 16,000 nuclei per sample were immediately incubated in a Transposition Mix to fragment DNA in open regions of chromatin and add adapter sequences to the ends of DNA fragments using the Chromium™ Single Cell ATAC Library & Gel Bead Kit (1000111). GEMs were generated by combining barcoded Gel Beads, transposed nuclei, a Master Mix, and Partitioning Oil on a Chromium Chip E (Product Code 1000086), and run using the Chromium Controller (10x Genomics). Libraries were generated with the Chromium Single Cell ATAC Library & Gel Bead Kit (1000111) and sequenced using the NextSeq 500/550 High Output Kit v2.5 (Illumina) on an Illumina NextSeq 550 platform. Using the 10x Cell Ranger ATAC 2.0.0 pipeline, scATAC-seq reads were de-multiplexed and aligned to the zebrafish genome (danRer11) to generate single cell accessibility counts. All data was deposited to GEO under accession GSE192947. Downstream bioinformatic analysis was performed in R using the packages Seurat (v4.0) and Signac (v1.4.0) (Butler et al., 2018; Stuart et al., 2019). We used the data integration algorithm implemented in Seurat to identify integration anchors and integrated all samples to remove batch effects. UMAP dimensional reduction was performed on the scaled data using Seurat function RunUMAP. Cell clusters were identified using Seurat function FindClusters. A gene activity matrix was generated from reads mapped to gene body and promoter (upstream 2 kb from the TSS), to facilitate cluster annotation by examining activity of cell-type-specific marker gene activity. Open chromatin peaks specific to cell clusters were identified using Seurat function FindAllMarkers. Differentially accessible peaks were calculated using Seurat function FindMarkers.

Using Signac, QC metrics for scATAC-seq experiments were computed by following the metrics outlined in the Analyzing PBMC scATAC-seq workflow (Stuart et al., 2019). The nucleosome binding pattern, TSS enrichment score, total number of fragments in peaks, and fraction of fragments in peaks were inspected. Cells that were outliers for QC metrics were removed. Datasets were normalized according to Signac TF-IDF normalization, feature selection was set to q0 for dimensional reduction, and linear dimensional reduction was run using SVD. Each LSI component and sequencing depth was correlated to assess if there was a strong correlation between the initial LSI component and the total number of cell counts. Cells were embedded in a low dimensional space and graph-based clustering and non-linear dimension reduction for visualization was performed according to Signac and Seurat. Chromatin accessibility associated with zebrafish genes was assessed. Gene coordinates were extracted and extended to include 2 kb upstream. The activities of canonical SC, PC, and HC marker genes were visualized. For integration with scRNA-seq, pre-processed scRNA-seq datasets were loaded using the methods for cross-modality integration and label transfer to confirm that scRNA-based classifications were consistent with the scATAC-seq data. scATAC-seq derived clusters were annotated.

Since Signac is an extension of Seurat, we used the FindAllMarkers function to obtain differentially expressed peaks for each of the clusters in a dataset with a min.pct of 0.025. Cluster assignments were made based on quantitative accessibility and gene expression. Using HOMER mergePeaks –d 100, we identified overlapping and unique peaks for each dataset in a cell type specific manner. To obtain RREs, common peaks shared between untreated and treated samples were removed. The peak data was used as input for downstream analysis including deep learning experiments.

### GREAT Analysis and Gene Ontology Annotations

GREAT version 3.0.0 (http://great.stanford.edu/great/public-3.0.0/html/) was used with the zebrafish assembly (danRer7, Jul/2010) (Hiller et al., 2013). Zebrafish enhancer genome coordinates were converted from danRer11 genomic coordinates to danRer7 coordinates for the analysis (liftOver; (Kent, 2002)). Functional annotations of zebrafish enhancers were performed with GREAT version 3.0.0 (Hiller et al., 2013) using the basal plus extension mode and default parameters (5 kb upstream, 1 kb downstream, plus distal up to 1000 kb). GO enrichment analysis was performed using AmiGO (Ashburner et al., 2000; Carbon et al., 2009; Gene Ontology, 2021).

### Motif analysis and deep learning co-enrichment analysis

To identify peak coordinates of interest containing instances of Sox and/or Six TF motifs, the annotatePeaks.pl function of HOMER with the “-m <MOTIF file<” option was used (Heinz et al., 2010). Motif file outputs by HOMER were selected based on initial *de novo* motif enrichment.

To differentiate accessible regions in the core promoter from those representing putative enhancers, we classified scATAC-seq peaks according to their position relative to the TSS of the closest gene. Thus, we classified regions that become more accessible during regeneration, and we looked for regions within a 50 kb genomic window. Peaks were annotated by genes (using bedtools window function) (Quinlan, 2014; Quinlan and Hall, 2010).

The overlapping probability was estimated by using the hypergeometric distribution using the phyper function in R: phyper(min(n1,n2),n1, n-n1,n2) - phyper(m-1,n1,n-n1,n2), where n is the total number of genes in the zebrafish genome, n1 is the number of genes in one list, n2 is the number of genes in the second list, and m is the number of overlapping elements in the given lists.

### Identification of co-occurrence of de novo motifs using deep learning (DL)

To discover de novo binding motifs, we built enhancer DL models based on ATAC-seq emerging peaks in progenitor cells, hair cells and support cells from regenerating inner ears (saccule and utricle). The positive set contained 18,845, 17,948, and 14,050 regulatory elements in each cell type, respectively. Each element was defined as a 400 bp region centered at the ATAC-seq emerging peak in the corresponding cell type. The control set contained 400 bp elements centered at the open chromatin regions from endothelial, liver and red blood cells in zebrafish (Ganis et al., 2012; Quillien et al., 2017). The training data set contained elements in positive set from all chromosomes, except for chromosomes 8 and 9, and 10-fold control sequences. The testing data set contained positive elements from chromosome 8 and 9 and 1-fold control sequences.

We constructed the DL model with four convolutional neural networks (CNNs) and detailed parameters can be found in FileS2. In the first CNN layer we used a sliding window of 9 bp (9-mer) and a step size of 3 bp to scan the input DNA sequences for de novo motifs, excluding the 50 bp regions from both ends to avoid potential errors near boundaries. We limited the maximum number of identified de novo motifs to 64 in order to reduce redundancy. For each de novo motif, we aligned all found 9-mer together and derived a position weight matrix (PWM). We next applied STAMP (Mahony and Benos, 2007) to find the best-matching known TF binding motifs for each derived PWM with a p-value < 1e-05.

To identify the possible co-occurrence between any pair of *de novo* motifs in a range of 100 bp in DNA sequences, we retrieved the frequency that both *de novo* motifs were detected in the same filter in the third CNN layers. If the frequency is significantly higher (p-value < 0.001) than background, the two *de novo* motifs were considered to be co-occurrences.

### CRISPR/Cas9 Enhancer Deletion

CRISPR/Cas9 mutagenesis was performed as previously described (Varshney et al., 2016). To study the effect of the upstream enhancer of *sox2* deletion on regeneration, we implemented the strategy described in Goudarzi et al. (Goudarzi et al., 2019). Cas9 protein was co-injected with 6 guide RNAs flanking the *sox2* upstream enhancer. Mutations rates were determined by PCR amplification using a pair of primers flanking the outermost guide RNA targets.

To identify the molecular lesion of the CRISPR-induced deletion, PCR products were amplified from DNA of F_1_ zebrafish containing heterozygous deletions, were then purified and subcloned into pCR2.1-TOPO vector by TA cloning. Following bacterial transformation, 4 colonies were picked and grown overnight. Plasmid DNA was extracted (Qiagen Miniprep Kit) and analyzed by Sanger sequencing.

6 sgRNA’s were designed to flank the upstream enhancer:

sox2-1955-l1 TAATACGACTCACTATAGGCAGGCGCACGAATTTAAAGTTTTAGAGCTAGAAATAGC sox2-1955-l2 TAATACGACTCACTATAGGCCGTCAATCATCACAAATGTTTTAGAGCTAGAAATAGC sox2-1955-l3 TAATACGACTCACTATAGGATACAATAAAGTAAACGCGTTTTAGAGCTAGAAATAGC sox2-1955-r1 TAATACGACTCACTATAGGCATTGTTTCTGGCACAGAGTTTTAGAGCTAGAAATAGC sox2-1955-r2 TAATACGACTCACTATAGGCCAACAAAAGCAGTCGTGGTTTTAGAGCTAGAAATAGC sox2-1955-r3 TAATACGACTCACTATAGGAGTCGTGAGGTAATGGGCGTTTTAGAGCTAGAAATAGC

### Hair cell and neuromast quantification

Hair cell staining and quantification were performed as described (Pei et al., 2016) using YO-PRO-1 (Life Technologies. Cat#: Y3603). For hair cell regeneration analysis, embryos from heterozygotic *sox2^hg138^* in-crosses at 5 dpf were treated with copper sulfate at 10 µM for 1 h at 28.5°C, recovered for 48 h at 28.5°C, and then counted for the regenerated hair cells in the lateral line neuromasts P1, P2, P4, and P5. Larvae were then genotyped to detect wild-type and enhancer deletion alleles. Approximately 40 embryos were used for each of the analyses except when otherwise indicated. The average number of hair cells and the standard error of the mean (s.e.m.) are shown in the graphs.

### Histological Methods

Adult zebrafish were euthanized using buffered MS-222. The heads were dissected and fixed in 4% formaldehyde overnight at 4°C. Inner ears were dissected as previously described in Liang and Burgess (Liang and Burgess, 2009).

### Hair Cell Labelling and Quantification

Alexa Fluor 488 phalloidin was used to visualize and quantify F-actin in stereocilia of zebrafish. Utricles and saccules were dissected and stained using Alexa Fluor 488 phalloidin as previously described in (Liang and Burgess, 2009; Liang et al., 2012). Proteins were detected in whole-mount utricles and saccules using standard immunofluorescence labeling methods. Primary and secondary antibodies used include the rabbit myosin VI and myosin VIIa antibodies (Proteus Biosciences 25-6790 & 25-6791, 1:300 dilution), rabbit Sox2 (Gentec #4477, 1:300-dilution), Alexa Fluor 568 goat anti-rabbit IgG (1:1000-dilution).

### Cellular Imaging and Analysis

Confocal images were acquired with a LSM 880 (Zeiss). Confocal Z stacks of the entire saccule and utricle were projected into a single image to capture all phalloidin positive cells from different planes of focus for counting. Phalloidin and Sox2 positive cells were counted with Image J/Fiji software (Schindelin et al., 2012) from the saccule and utricle whole mounts.

## Supplementary Files

Table S1. Differentially expressed genes between cell types of larval lateral line neuromast and adult inner ear, Related to Figure S1.

Table S2. Cluster marker genes enriched in non-regenerating saccule and utricle, Related to Figure S2A.

Table S3. Differentially expressed genes between individual clusters from non-regenerating saccule and utricle, Related to Figure S2A.

Table S4. Cluster marker genes between clusters of regenerating saccule and utricle, Related to Figure S2B

Table S5. Differentially expressed genes between clusters of regenerating saccule and utricle. Related to Figure S2B

Table S6. Cluster marker genes of 12 sample scRNA-seq aggregate, Related to Figure 2

Table 7. Cluster marker genes and differentially expressed genes enriched during hair cell regeneration, Related to Figure 3

Table S8. Gene module data generated using Monocle3, Related to Figure 4.

Table S9. Single-cell sample library statistics, Related to Figure 5

Table S10. scATAC-seq cluster markers generated using Signac and Seurat, Related to Figure 5

Table S11. RREs identified in support cells, progenitor cells, and hair cells used for downstream analysis, Related to Figure 6

Table S12. Genes +/- 50kb of RRE in support cells, progenitor cells, and hair cells, Related to Figure 6

Table S13. Genes +/- 50kb of RRE and CNE in support cells, progenitor cells, and hair cells, Related Figure 6

Table S14. Differentially expressed genes identified during hair cell regeneration including differential genes +/- 50kb of RRE (and CNE), Related to Figure 6

Table S15. Hypergeometric distribution tests, Related to Figure 6

Table S16. Summary of differentially expressed genes linked to RREs containing predicted binding TF sites, Related to Figure 7

File S1. Convolutional layers and parameters used in supervised deep learning models, Related to Figure 7

**Figure S1.**
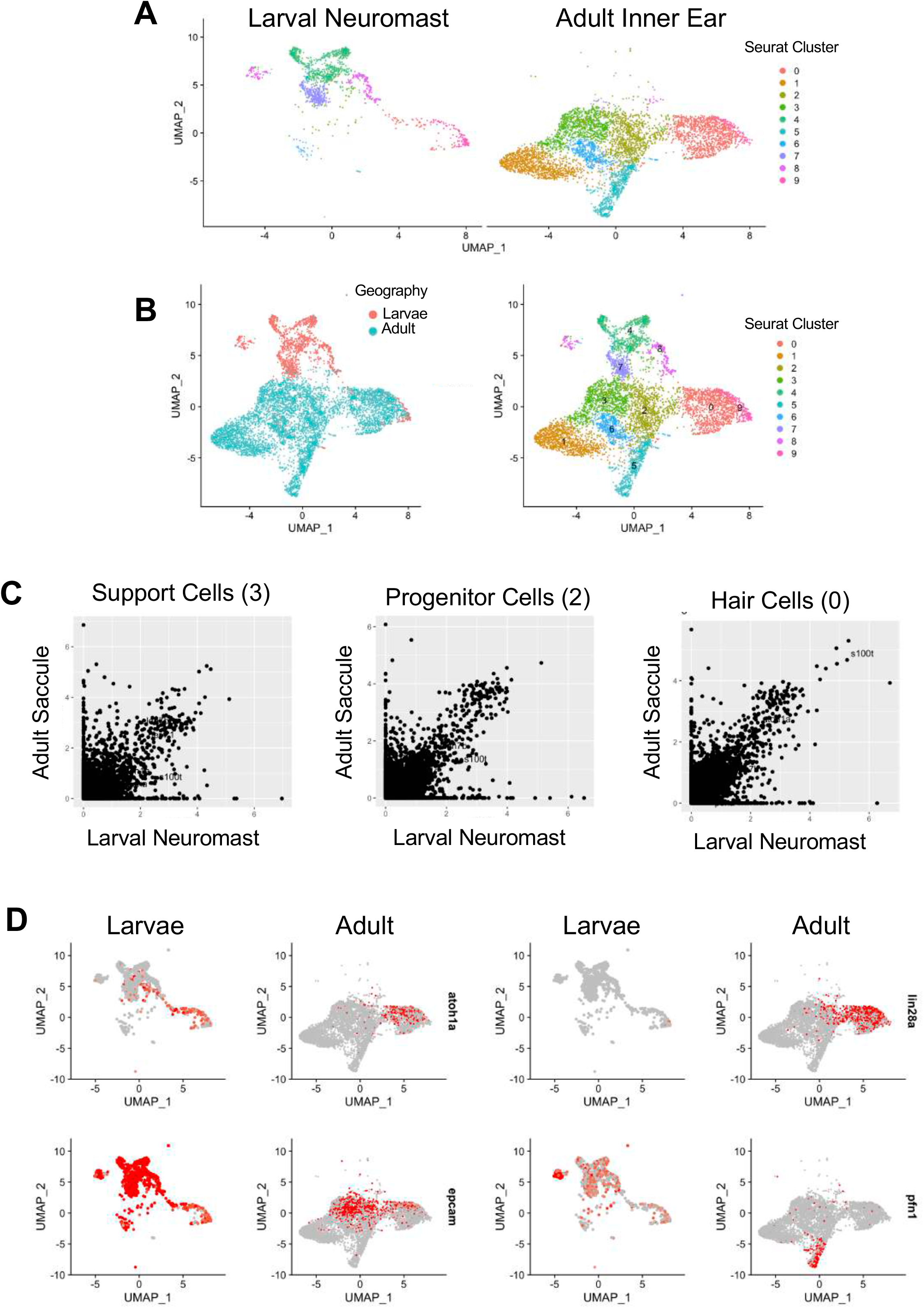
Adult inner ear cells are transcriptionally distinct from larval lateral line neuromast cells. (A) Unbiased clustering of larval neuromast (left) and adult inner ear tissue (right). (B) Geography of clustered larval neuromast and adult inner ear. (C) Correlation plots comparing cell types. (D) Gene expression profiles of select genes.

**Figure S2.**
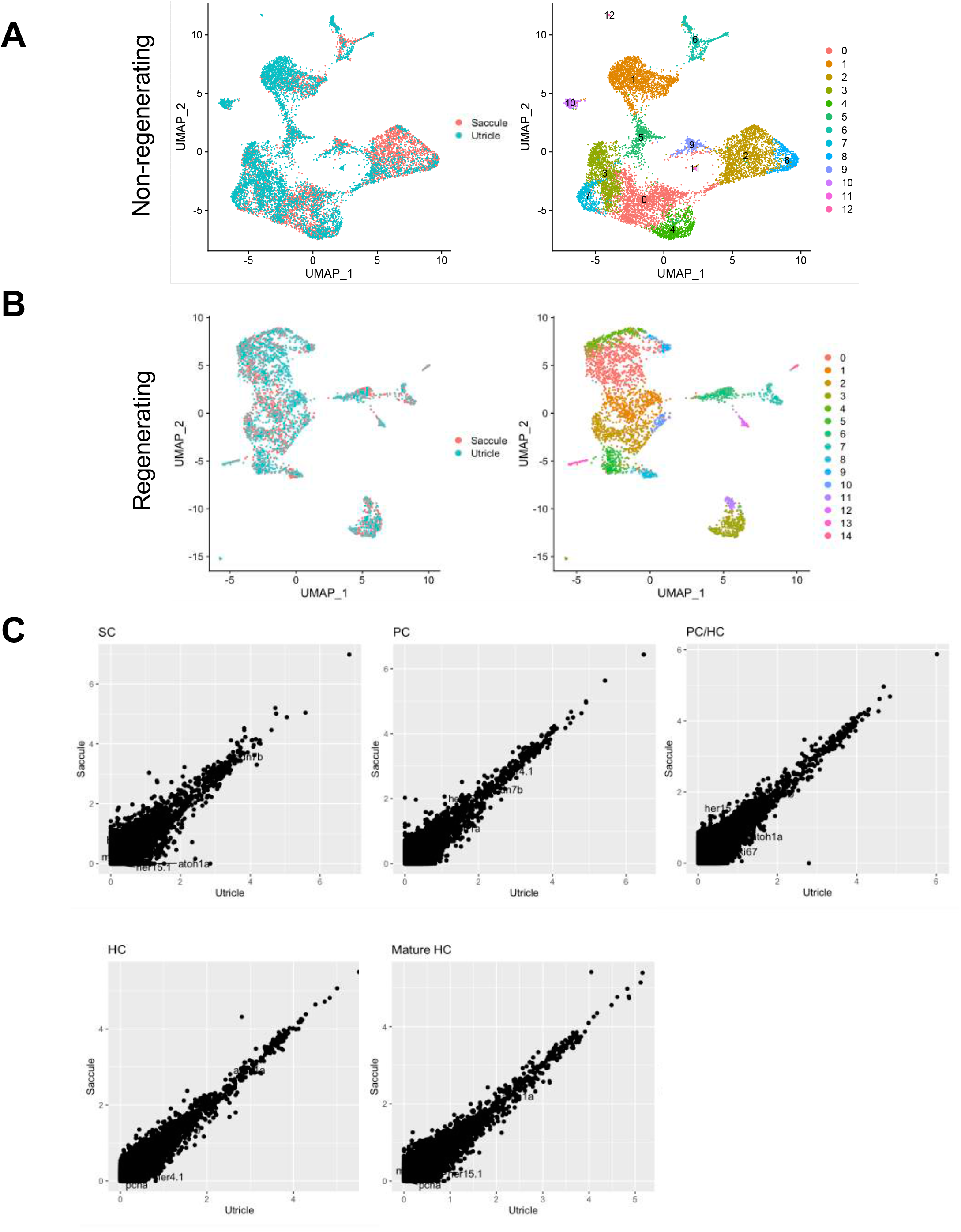
Organ-level similarities between complementary cell types. (A) Non-regenerating saccule and utricle overlayed (left) and clustered (right). (B) Regenerating saccule and utricle overlayed (left) and clustered (right). (C) Scatter plot analysis of differentially expressed genes in sensory cell types during regeneration. Demonstrates that the organs are highly similar in gene expression.

**Figure S3.**
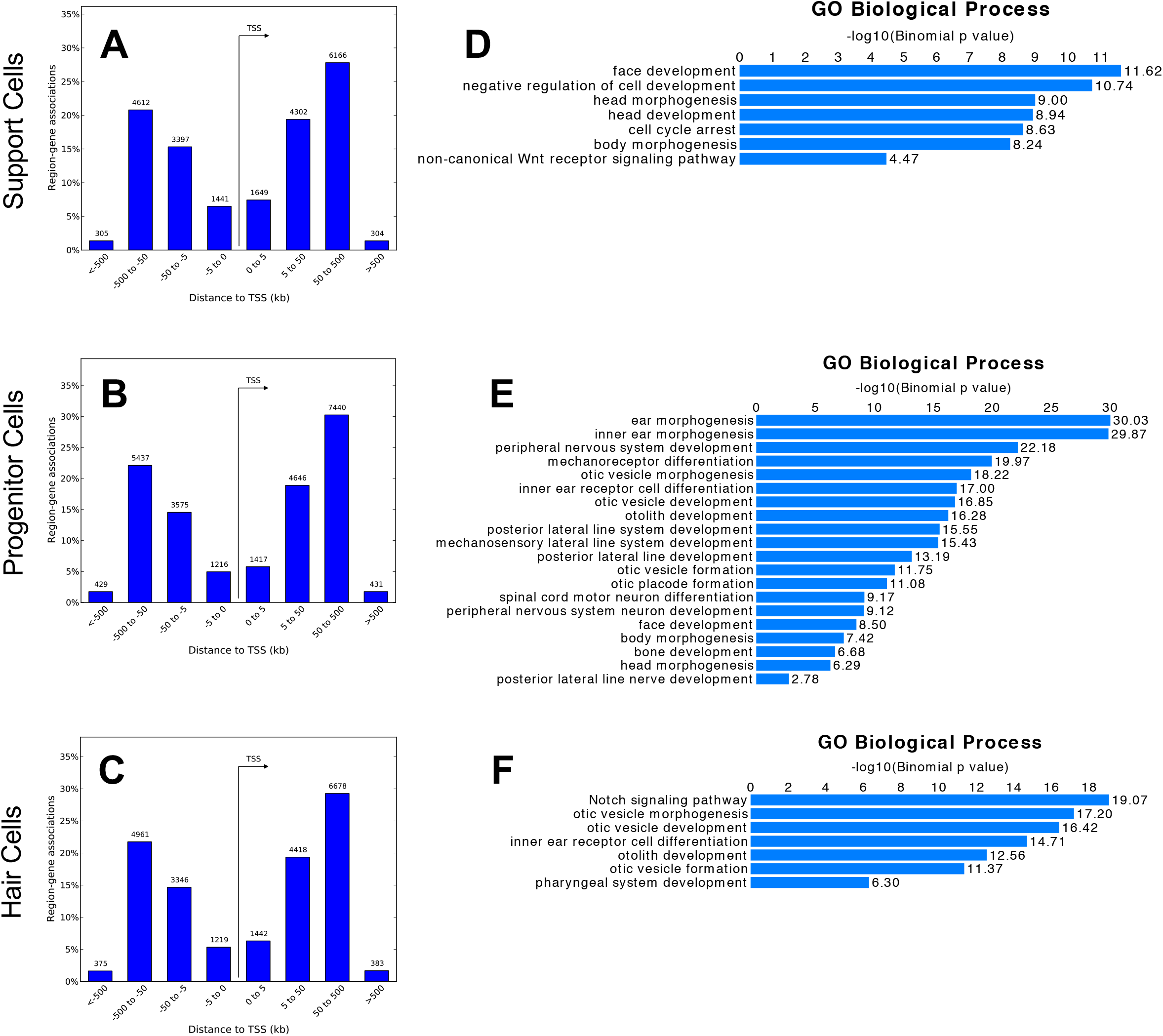
GREAT analysis on emerging peaks in regenerating inner ear tissues. (A-C) Distance from predicted peak to gene TSS. (D-F) GREAT GO biological processes in predicted cell-specific enhancers.

**Figure S4.**
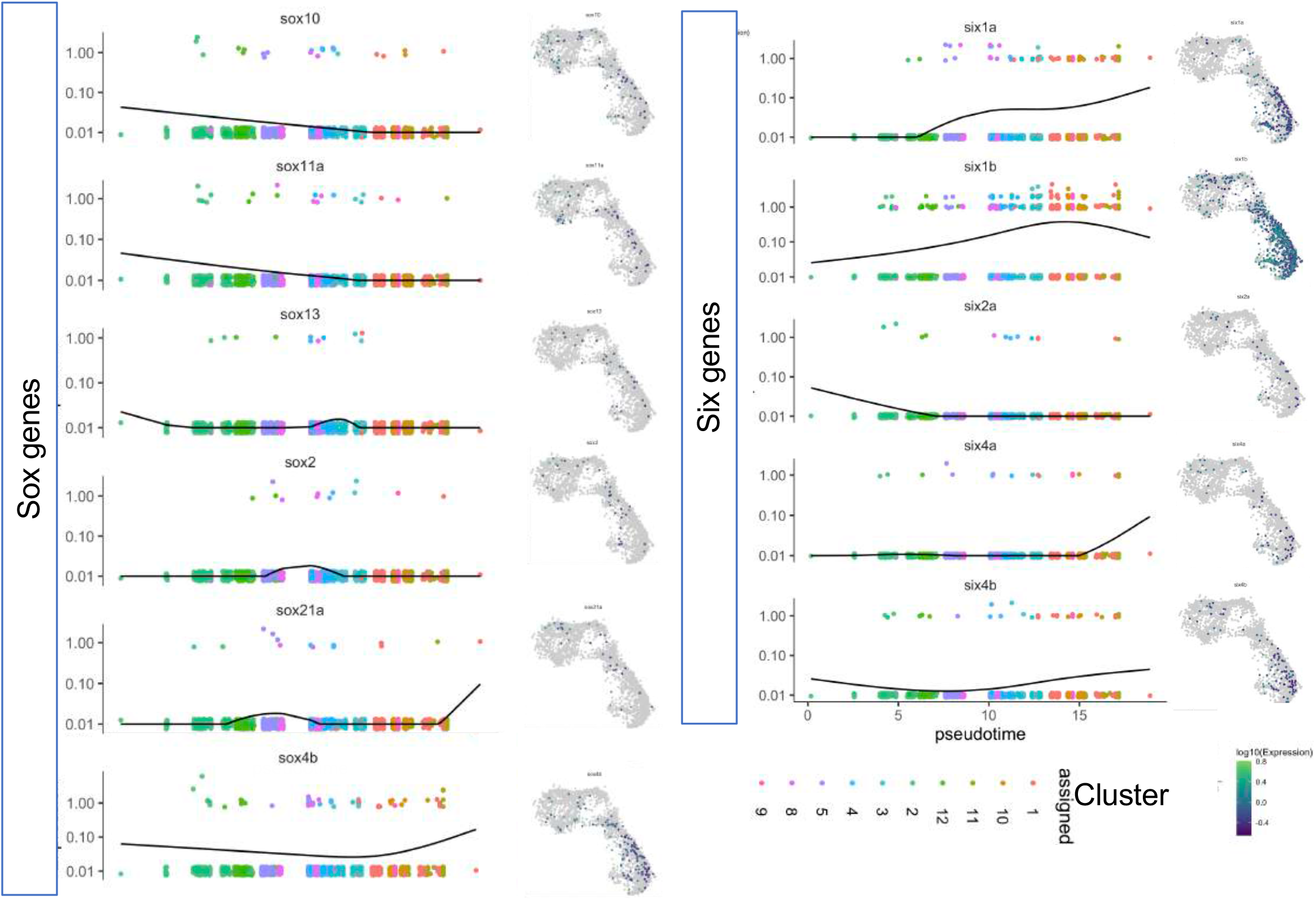
*sox* and *six* gene expression alterations during hair cell regeneration. Pseudotime ordered single cell expression trajectories for all Sox (left) and Six (right) genes with detectable differential expression.

**Figure S5.**
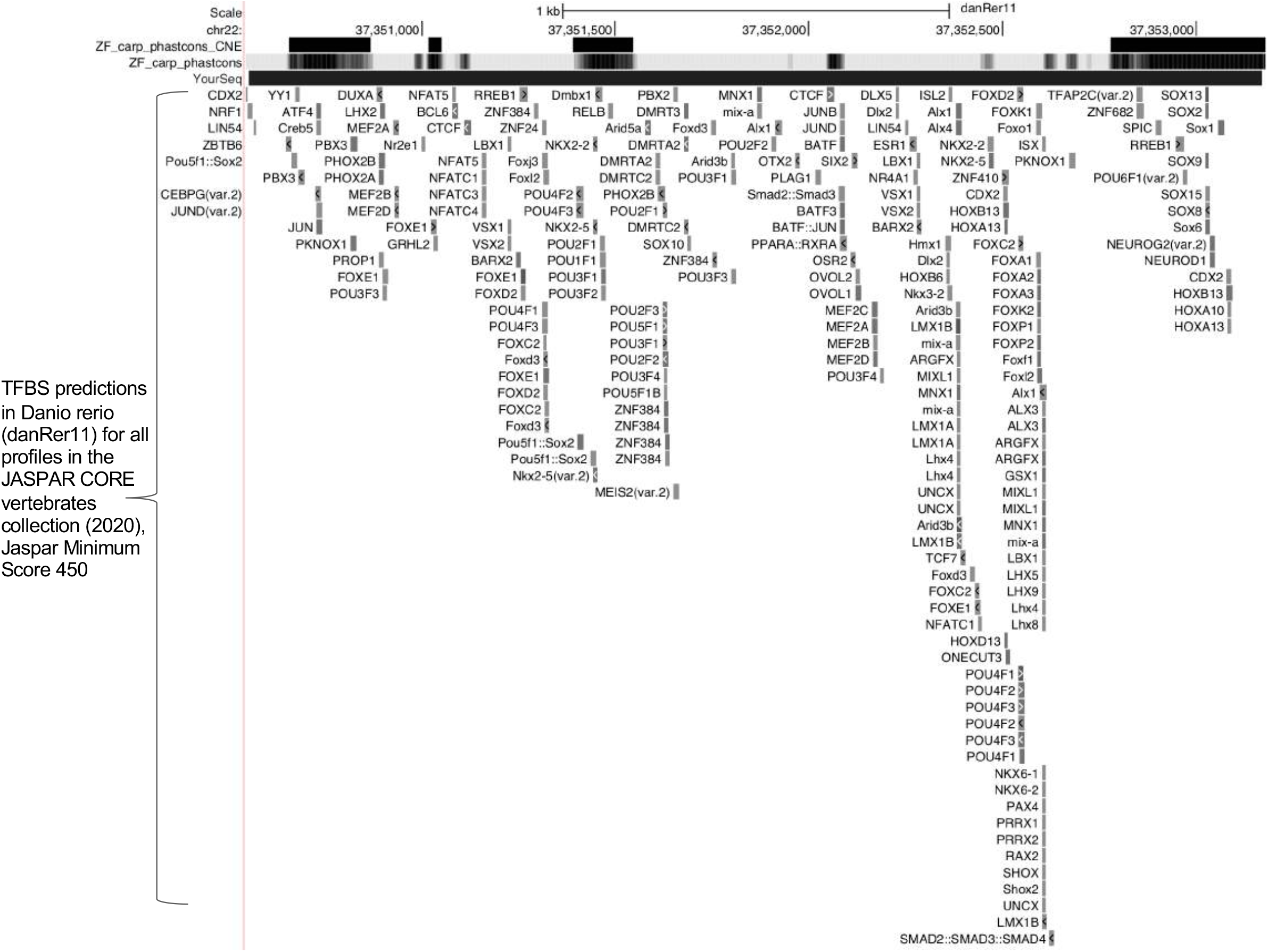
*sox2^hg138^* region. UCSC Genome Browser screen shot of the deleted region. Custom tracks on danRer11 in the UCSC Genome Browser showing the zebrafish/goldfish/carp alignments (ZF_carp_phastcons_CNE and ZF_carp_phastcons). Black bar denotes deleted region (YourSeq). TFBS predictions in Danio rerio (danRer11) for all profiles in the JASPAR CORE vertebrates collection (2020) are in grey (Minimum score 450).

**Figure S6.**
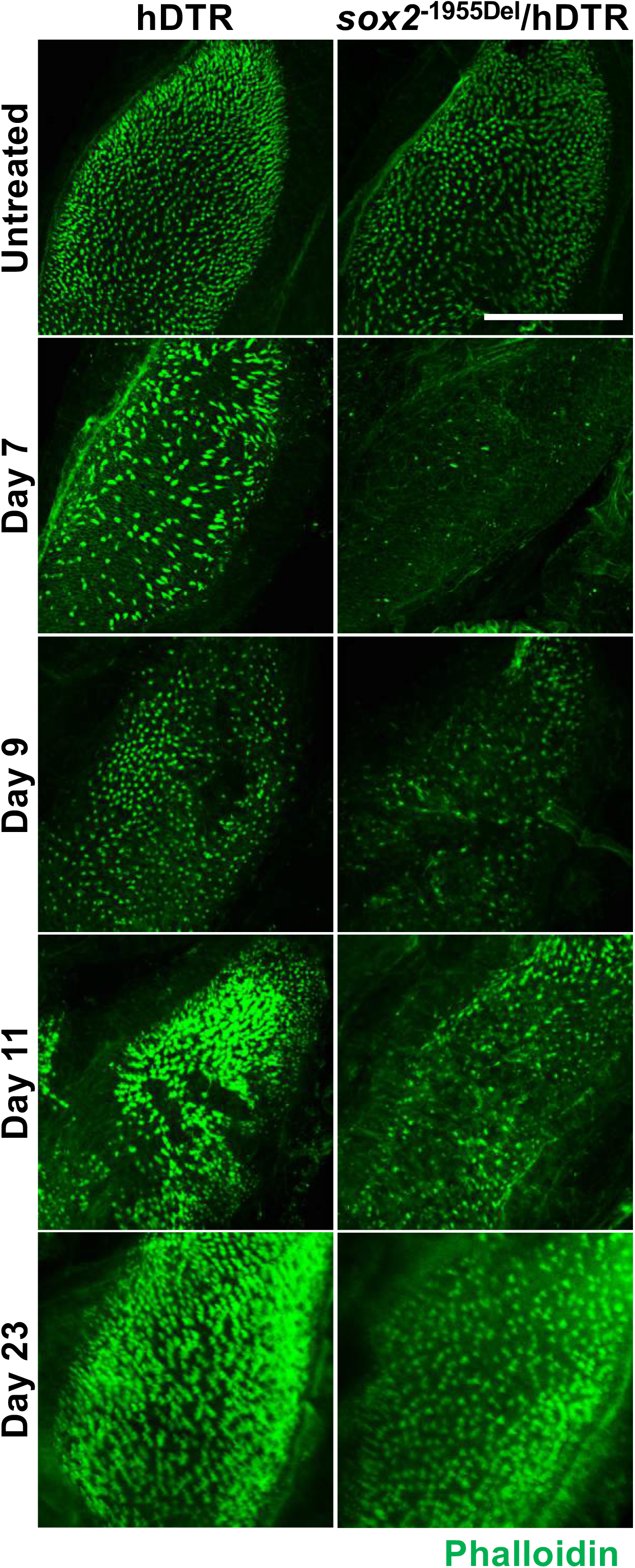
*sox2^hg138^* enhancer deletion mutants exhibit severely delayed hair cell regeneration in auditory sensory epithelia. Saccule isolated from Tg(*myo6b*:hDTR) and *sox2^hg138^* zebrafish following DT injection on days 7, 9, 11, and 23. Phalloidin staining (green) labels hair cell stereocilia. Brightness and contrast adjusted to 30% and 40%, respectively. Scale bar 100 µm.

